# Ferroptosis susceptibility of melanoma cells: dependence on cell-type, acquired drug resistance, and medium composition

**DOI:** 10.1101/2024.08.12.607529

**Authors:** Jasmin Renate Preis, Catherine Rolvering, Mélanie Kirchmeyer, Dzeneta Halilovic, Joanna Patrycja Wroblewska, Fabrice Tolle, Iris Behrmann, Claude Haan

## Abstract

Resistance of melanoma cells to targeted therapy (BRAF and MEK inhibitors) is a major clinical problem and alternative treatments are sought. We describe the establishment of modular physiologic medium (MPM) and Mel-MPM (which contains additional supplements and sustains the 3D growth of melanoma cells, fibroblasts (NHDFs) and HMEC-1 endothelial cells) as novel resources for melanoma and combine them with a multi-cell-type matrix-embedded 3D culture model to investigate melanoma cell vulnerabilities in a more physiological setting.

We made use of the modular nature of MPM to interrogate NEAA dependencies in melanoma cells and we found them to be particularly sensitive to the depletion of C/C. We additionally describe that melanoma cells are less sensitive to ferroptosis inducing compounds when cultured in MPM compared to RPMI and we could attribute this to different components of MPM and Mel-MPM (selenite, B27). Cell death induced by the glutathione peroxidase 4 inhibitor, ML162, had characteristics of ferroptosis or apoptosis depending on cell type, its drug resistance status and the culture medium. Cystine/cysteine starvation and ML162 treatment combinations increased melanoma cell death in 2D, 3D, and also in the complex matrix embedded multi-cell-type-3D system in Organoplates^TM^. This underlines the potential of combining metabolism-oriented drug treatments with amino acid starvation conditions, which is of interest in view of future therapeutic approaches to combat melanoma and other cancer types.

## Introduction

Melanoma, an aggressive form of skin cancer (Whiteman et al., 2016), is often associated with mutations in *BRAF* (e.g. V600E) which are responsible for abnormal MAPK pathway signaling and affect cellular metabolism (Alkaraki et al., 2021; Avagliano et al., 2020). Treatments with the BRAF inhibitor, Encorafenib, and the MEK inhibitor, Binimetinib, have increased the 5-year survival rate to 35% (Dummer et al., 2022). However, many patients do not respond to therapy or rapidly develop resistance to MAPK pathway inhibitors and immune checkpoint blockade treatments (Winder & Virós, 2018), highlighting the urgent need for new therapeutic strategies. A number of mechanisms conferring kinase inhibitor resistance have been described (Kozar et al., 2019). Melanoma cells show a high metabolic plasticity, with drug resistance being linked to metabolic rewiring (Alkaraki et al., 2021; Avagliano et al., 2020; Pendleton et al., 2023).

The tumour heterogeneity and the microenvironment influence many features of the cellular metabolism of cancer cells. Inhibitory compounds identified in preclinical *in vitro* models (standard 2D cell culture models) regularly perform differently in complex cell models or mouse models (Muir et al., 2018) and thus often fail to enter clinical trials. Importantly, studies using 3D cell culture models have been reported to predict the tumour drug response more accurately (Lee et al., 2018; Vlachogiannis et al., 2018). Recently more physiologic media formulations have been described which contain metabolites at physiological concentrations (Cantor et al., 2017; Vande Voorde et al., 2019). The human plasma metabolome is estimated to contain well over 4000 metabolites (Psychogios et al., 2011); this is of course not fully reflected by these media, but they contain the most abundant soluble polar metabolites found in human plasma. Striking differences in cellular metabolism and drug action have been observed in these media compared to standard cell culture media (Cantor et al., 2017; Rossiter et al., 2021; Vande Voorde et al., 2019).

Our aim was to culture our drug-sensitive and drug-resistant cell lines in more physiologic conditions for further drug screening and/or -validation approaches and to identify treatments which target especially drug-resistant melanoma cells. We therefore established more physiologic media (MPM and Mel-MPM) to better recapitulate the physiological metabolite concentration. We first depleted a number of non-essential amino acid, individually or in pairs, to uncover possible vulnerabilities of the drug-sensitive and drug-resistant melanoma cells in MPM. Cystine/cysteine withdrawal as well as treatment with ferroptosis inducing compounds (ML162, Erastin) resulted in much less ferroptosis in physiological medium compared to RPMI. Drug-(Encorafenib/Binimetinib) resistant cell lines were generally more vulnerable to amino acid withdrawal. Induction of either ferroptosis or apoptosis depended on the cell line studied and the culture medium used. Focussing on ML162, we explored different combination treatments (starvation or drug treatment) to increase either the ferroptosis or apoptosis induction we observed. Finally, we validated these results in a matrix-embedded multi-cell-type 3D cell culture system.

## Materials and methods

### Standard cell culture and materials

All cells were grown at 37°C in a water-saturated atmosphere at 5% CO_2_. All melanoma cell lines (624Mel (Dr. Ruth Halaban), A375 (ATCC, cat.no. CRL-1619), WM3248 (Rockland, cat.no. WM3248-01-0001), MelJuso (DSMZ, cat.no. ACC 74) and M45 (Dr. Dagmar Kulms) were maintained in RPMI (Gibco) supplemented with 10% FBS. Before experiments, the melanoma cells were sub-cultured for 1 passage in MPM if not indicated differently. HMEC-1 cells (ATCC) were maintained in MCDB131 (Gibco) supplemented with 10 ng/mL Epidermal Growth Factor (EGF, Peprotech), 1 µg/mL Hydrocortisone (Merck), 10 mM Glutamine (Gibco), and 10% FBS (Gibco). NHDF cells (PromoCell) were cultivated in DMEM (Gibco) supplemented with 25 mM HEPES (Gibco), and 10% FBS (Gibco). Adaptation of NHDF and HMEC-1 to Mel-MPM was performed 7-10 days before starting the experiments. HMEC-1 cells were transduced with pLV-Bsd-CMV-tagBFP (Bio-connect) and selected with 10 µg/mL of blasticidin to yield the HMEC1-BFP cells. NHDF-iRFP cells were generated by transduction with LV-iRFP713-P2A-Puro (Imanis), selected with 1 µg/mL puromycin and sorted on a FACS Aria (BD biosciences).

### Generation of drug-resistant (DR) melanoma cell lines

The drug-resistant melanoma cell lines (624Mel-DR, A375-DR, WM3248-DR) were generated by prolonged exposure with the BRAF inhibitor Encorafenib and the MEK inhibitor Binimetinib (MedKoo biosciences). Briefly, the concentrations used to generate drug-resistant (DR) cells (624Mel: 30 nM Encorafenib, 400 nM Binimetinib; WM3248: 15 nM Encorafenib, 200 nM Binimetinib), correspond approximatively to 10-times the IC50 concentrations of both drugs individually. The exposure was continued until the cells re-proliferated in the presence of drug. Subsequently, the resistant cell lines were maintained under continuous exposure to Encorafenib and Binimetinib. WM3248-DR and 624Mel-DR cells were transduced with firefly luciferase-GFP lentivirus (CMV, Puro; Cellomics) and selected with 0.5 or 0.75 µg/mL puromycin to yield the WM3248-DR-GFP/luc and 624Mel-DR-GFP/luc cells, respectively. Because we do not describe luciferase assays in this publication, the cells are referred to as WM3248-DR-GFP and 624Mel-DR-GFP in the following text. GFP-positive cells were sorted on a FACS Aria (BD biosciences) before amplification and freezing.

### Modular physiologic medium (MPM) preparation

The preparation protocol is based on a protocol published before (Vande Voorde et al., 2019) with some adjustments introduced concerning concentrations and number of component solutions. The modular physiological medium (MPM) is built from 9 metabolite component solutions, the BME vitamin mix, and dialysed FBS, all of which were stored at −80°C (see Table S1). Solutions 1, 2, 3, 4, 5, and 9 are stored as sterile filtered 2 mL aliquots (250X solutions; 2mL/ 500mL MPM). Solution 6 to 8 are stored as sterile filtered 1 mL aliquots (500X solutions; 1mL/ 500mL MPM). The BME vitamin mix (Merck) was supplemented with Vitamin C and Vitamin B12 and used as 100X solution (5 mL/500 mL MPM). The FBS concentration in the medium was 2.5%. FBS (Gibco) was dialysed using the SpectraFlow Lab system (Repligen) and SnakeSkin dialysis tubing (3.5K MWCO, 35 mm; Thermo Fisher Scientific PI88244) for 36 hours, so that the metabolite concentration of FBS was reduced by a factor of around 100-fold, while proteins and substances with a mass higher than 3.5 kDa were retained. We also added 25 mM HEPES to stabilize the pH for microscopy experiments. The sterile solutions 1 to 9 were added with sterile tips, while 12.5 mL of dialysed FBS, 12.5 mL HEPES (1 M, pH 7.4), and 5 mL Vitamin mix were sterile filtered upon addition to the EBSS. EBSS (Merck) basal medium was used at 0.9x of its original concentration to ensure that the total concentration of the MPM medium (151 mM) was near the physiological concentration of 150 mM. The osmolality of MPM, measured using an ARKRAY-Osmostation OM6050 was ∼ 290 mOsmol/kg, which was within the normal reference range for human plasma (275-299 mOsmol/kg). For the experiments with MPM lacking a specific non-essential amino acid (NEAA), we prepared individual solutions of the NEAA and added them to MPM lacking either solution 2 or solution 3, depending on which amino acids were investigated.

### Mel-MPM (Melanoma-MPM)

To better mirror physiological conditions, promote growth of 3D spheroids, and cultivate melanoma cells with fibroblasts and endothelial cells embedded in matrix, we established Mel-MPM. MPM was supplemented with xeno-free B27 supplement (Gibco) and with growth factors (Insulin 500 ng/mL, HGF 50 ng/mL, BMP4 20 ng/mL, EGF 5 ng/mL, Hydrocortisone 1 μg/mL). Mel-MPM was used for growth of melanoma lines, NHDF fibroblasts and HMEC1 endothelial cells as well as of mixed cultures of NHDF, HMEC1, and melanoma cells in 3D culture.

### Vascular tube formation assay

HMEC1 cells were seeded on Matrigel^®^ Growth Factor Reduced (GFR) Basement Membrane Matrix, Phenol Red-free, LDEV-free (Corning), at a density of 24,000 cells per 96-well in either MCDB131 medium (with EGF and hydrocortisone) or Mel-MPM. 23 hours after seeding, cells were stained with 8 µM Calcein AM (live cell stain; Cayman Chemical) and 1 µg/mL Hoechst 33342 (Live cell nuclear stain; Life Technologies) for 1 hour and imaged on a Cytation 5 imaging reader (Agilent Technologies) using DAPI and GFP filter/LED cubes. One representative biological replicate out of three is shown.

### Immunofluorescence microscopy and confocal imaging

NHDF cells were seeded in DMEM or Mel-MPM on 8-well µ-slides (Ibidi), fixed with 4% PFA and treated with 0.1% Triton-X100. After overnight incubation with primary antibodies (mouse anti-Fibroblasts Antibody, clone TE-7 (Merck, Cat# CBL271, RRID: AB_93449, diluted 1:100) or rabbit anti-α-Smooth Muscle Actin XP (Cell Signaling, Cat# 19245S, RRID: AB_2734735, diluted 1:500) and subsequent incubation with secondary antibodies (donkey anti-mouse Alexa 488 or donkey anti-rabbit Alexa 647, both 1:500) and DAPI (1 µM), cells were imaged on a Cytation 5 imaging reader (Agilent Technologies) using DAPI, GFP, and Cy5 filter/LED cubes. One representative biological replicate out of three is shown.

For live cell confocal imaging, the live cell stains Hoechst 33342 (Life Technologies), MitoLite Red FX600 (AAT Bioquest), and MitoSox Green (Life Technologies) were used at the concentrations indicated in the Figure legends. Cells were stained in MPM without FCS for 1h. Thereafter, medium was replaced with full MPM and cells were incubated with or without 5 ML162 for 50min. Confocal imaging was performed on a Cytation 10 imager (Agilent technologies) using DAPI, GFP, and TRITC filter cubes at 37°C and 5% CO_2_. Image analysis was carried out using Gen5 3.14 (Agilent technologies). Mitolite Red FX600 was used to create masks inside which the MitoSox Green intensity was quantified. Graphs were plotted using Graphpad Prism 10.2.2. One representative biological replicate out of three is shown. For each biological replicate, 6 to 8 images were analysed for each condition, mean and standard deviation are shown.

### 2D cell proliferation and live/dead assays

Cells were seeded in 96-well flat-bottom plates (Greiner Bio-one) for 2D adherent cell growth. Treatments were applied as described in the Figure legends. After 3 or 4 days, cells were stained with Hoechst 33342 (1 µg/mL, nuclear live cell stain, Life Technologies) and Sytox Orange (1 µM; nuclear dead cell stain, Life Technologies) for at least 30min and imaged on the Cytation 5 instrument (Agilent Technologies) using DAPI and RFP filter/LED cubes. The Gen5 software (Agilent) was used to determine total cell numbers (using Hoechst-stained nuclei) and cell death (Sytox orange-positive nuclei). Alternatively, cell growth was measured using the CyQuant Direct Cell Proliferation Assay Kit and PrestoBlue (both Life Technologies) in a parallel detection protocol. Fluorescence of CyQuant and PrestoBlue was measured on the Cytation 5 instrument (CyQuant: λex: 489 ± 12nm / λem: 530 ± 10nm; PrestoBlue: λex: 547 ± 12nm / λem: 612 ± 20nm). The graphs were plotted using Graphpad Prism 9. One representative biological replicate out of three is shown. Each biological replicate was performed in three technical replicates, and the mean and standard deviation are plotted. IC50 values were determined from biological replicates (n=3) using GraphPad Prism 9, log [inhibitor] *vs*. response-variable slope 4PL curve fit.

### Caspase activity assay

Cells were seeded in 96-well black μclear plates (Greiner Bio-One) one day before being incubated for 16h C with or without ML162 (5 μM) or Staurosporine (2 μM, positive control) treatment. To monitor the amount of background apoptosis in the cultures, 5 μM of the caspase inhibitor Ac-DEVD-CHO was added to one of the two untreated sample wells just before lysis. The cells were lysed by adding 50 μL of a 3 times concentrated Reaction and Lysis (RELY) Buffer (containing 150 mM Tris pH7.4, 300mM NaCl, 30% glycerol, 1% Triton, 0.3% Chaps, 6mM EDTA, 4 mM DTT, and 75 μM of the caspase substrate Ac-DEVD-AFC) onto the 100 μL cell culture volume per 96 well. After 30min to 1h incubation at room temperature, the AFC (7-amino-4-trifluoromethyl coumarin) fluorescent signals were measured on a Cytation 5 instrument (Agilent Technologies) using the fluorometer function (410-20nm excitation bandwidth, 490-520nm detection bandwidth). One representative biological replicate out of three is shown. Each biological replicate was performed in three technical replicates, and the mean and standard deviation are shown.

### 3D cell viability and cell death assays

624Mel-DR-GFP and WM3248-DR-GFP cells were seeded in 96-well ultralow attachment U-bottom plates (FaCellitate) in MPM or MPM with reduced amino acid levels for spheroid growth. The culture medium of spheroids was replaced after 3 to 4 days of culture. To track spheroid growth over time, brightfield images of the spheroids were taken 1 day after seeding and then approx. every 2^nd^ day with a Cytation 5 instrument (Agilent Technologies). On day 7, cells were stained with Propidium Iodide (PI, 4 µg/mL) for 3h to check for cell death within the spheroid. Sphere size/volume, GFP, and PI were measured on the Cytation 5 instrument using the LED/filter cubes for GFP and PI. Image processing was performed using the Agilent BioTek Gen5 software. The brightfield image was used to create a mask around the sphere; GFP and PI mean intensities were calculated within this mask. The average size (diameter) of round spheres was used to calculate the volume. One representative biological replicate out of 3 is shown. For each biological replicate, six technical replicates were analysed for each condition, and the mean and standard deviation are shown.

### Lipid peroxidation assay

After 20h of AA starvation or after 2h of drug treatment, cells were incubated with 2 µM BODIPY 581/591 C11 (Thermo Fisher Scientific) for 30min. Cells were washed with PBS, trypsinized, and resuspended in fresh medium before fluorescence-activated flow cytometry analysis (excitation 488nm). Lipid peroxidation was recorded for 20,000 events (single cells) in the FITC channel. As a positive control for lipid peroxidation, cells were treated with 50-200 µM cumene hydroperoxide (CHP) for 2h and analyzed by flow cytometry. One biological replicate out of three is shown, except for A375-DR (only two biological replicates were performed).

### Cell lysis and Western blot immunodetection

Cell lysis and Western blot analysis was performed as described before (Vollmer et al., 2009). Proteins were separated using self-cast Tris/acetate gradient gels, followed by electro-blotting onto a PVDF membrane (PVDF-PSQ or PVDF-FL, Millipore). The following antibodies were used for detection at a dilution of 1:1000: cPARP antibody (Cell Signaling Technology Cat# 5625, RRID: AB_10699459), eIF2α (Cell signaling; Cat# 5324, RRID: AB_10692650), GAPDH (Sigma; Cat# G9545, RRID: AB_796208), STAT1 (BD biosciences, Cat# 610116, RRID: AB_397522), NRF2 (ThermoFisher Scientific; Cat# PA5-27882, RRID: AB_397522). HRP-conjugated secondary antibodies were purchased from Cell Signaling. Signals were detected using a self-made ECL solution (containing 2.5 mM luminol, 2.6 mM hydrogen peroxide, 100 mM Tris/HCl pH8.8, and 0.2 mM para-coumaric acid) (Haan & Behrmann, 2007) or the SuperSignal™ West Femto Maximum Sensitivity Substrate (Thermo Fischer Scientific). The ECL signals were captured using a CCD camera system (Fusion FX, Vilber). One representative biological replicate out of three is shown.

### Cell culture in OrganoPlates

For seeding in the OrganoPlates™ (Mimetas), cells were counted, pelleted and embedded in Geltrex LDEV-free reduced growth factor basement membrane matrix (Gibco, final concentration around 11 mg/mL). 1×10^4^ WM3248-DR-GFP, 8×10^3^ HMEC-1-BFP cells, and 5×10^3^ NHDF-iRFP cells were embedded per µl of Geltrex. 2 µL of the cell-matrix mix were added to each gel inlet channel of a 3lane40 OrganoPlate™ (Mimetas). After 15min at 37°C to allow the gel to solidify, Mel-MPM with 2.5% FBS was added to the medium channels. After 2 days of pre-incubation, the medium was replaced with fresh Mel-MPM with 2.5% FBS or with modified media (e.g. Mel-MPM without cysteine (Mel-MPM-C) or medium containing drugs, which was also refreshed after 2 days). OrganoPlates™ were kept on an OrganoFlow shaker in the incubator to allow perfusion (Mimetas; 7° inclination, shaking interval time 8min). 4 days after drug addition, the cells were imaged on a Cytation 10 instrument (Agilent Technologies). Mean fluorescence intensities inside the matrix channel were quantified using Gen5 3.14 (Agilent Technologies) and plotted using Graphpad Prism 10.2.2. One representative biological replicate out of three is shown. Each biological replicate was performed in 4 technical replicates.

## Results

### The modular physiologic medium MPM consistently supports growth of melanoma cells in 2D and 3D cultures

Traditional cell culture media differ substantially from human plasma or interstitial fluid to which cancer cells are exposed (Ackermann & Tardito, 2019). We aimed to implement a modular physiological medium (MPM) resembling human plasma, based on previously published metabolite concentrations in human plasma (Dereziński et al., 2017; Psychogios et al., 2011; Trabado et al., 2017; Wang et al., 2018) and especially based on two recently established physiologic media formulations (Cantor et al., 2017; Vande Voorde et al., 2019). In short, we included those metabolites that were described in at least two studies, and which were measured at a concentration ≥ 6 μM. MPM contains 26 additional metabolites not present in most standard media, which account for 6.7 mM (45%) of the total nutrient concentrations in MPM and is thereby similar to the physiologic media HPLM and Plasmax (Cantor et al., 2017; Vande Voorde et al., 2019) (see Figures 1A, S1A/B and Table S2). Upon culturing our melanoma cell lines in MPM, we observed slight phenotypic changes, but all cells proliferated well even though we used only 2.5% FCS (to reduce the amount of FCS along the 3R guidelines) (data not shown). Our observations recapitulate studies by others who also observed phenotype changes but consistent growth of the cells in physiologic media (Avellino et al., 2023; Cantor et al., 2017; Golikov et al., 2022; Torres-Quesada et al., 2022; Vande Voorde et al., 2019).

**Figure 1.**
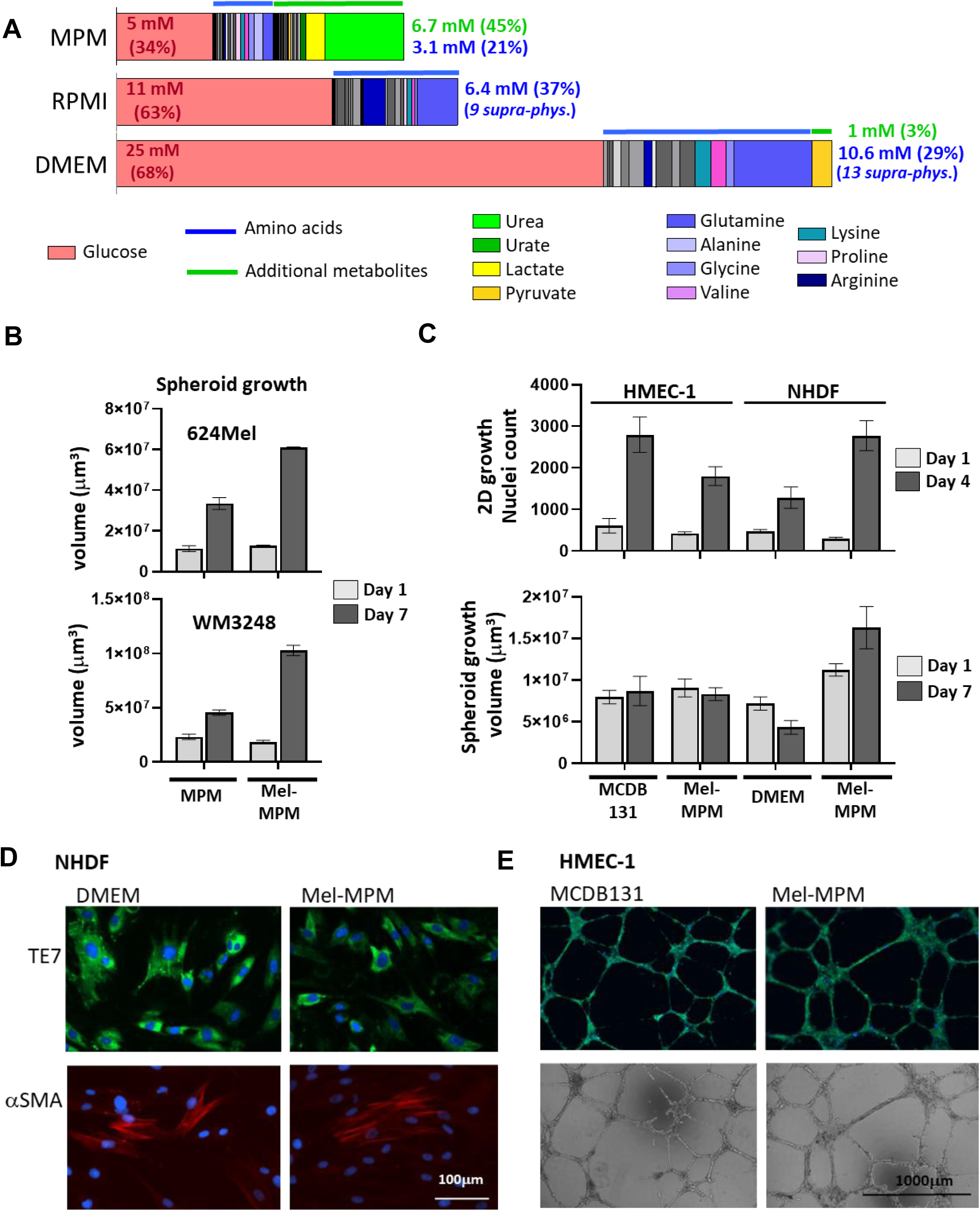
The physiologic media MPM and Mel-MPM and their effects on model cell cultures. A) The bar diagrams indicate the composition of main nutrients (glucose, amino acids and other metabolites) of the physiologic MPM and the two classic culture media used in this study, RPMI1640 and DMEM. The total concentrations and the percentages of glucose (red), amino acids (blue lines and writing) and other metabolites (green lines and writing) are indicated. B) Spheres of 624Mel and WM3248 cells were grown in MPM and Mel-MPM for 7 days, the volumes of the spheres are plotted. To illustrate the growth, the volumes of the spheres at day 1 and day 7 are shown. C) 2D and spheroid growth (measured microscopically by nuclei count or sphere volume determination, respectively) of HMEC-1 and NHDF cells in their standard media (DMEM and MCDB131, respectively), compared to Mel-MPM. 2D growth was followed for 4 days, while 3D spheroids were grown for 7 days. D) Immunofluorescence stain of 2D cultures of NHDF with TE7 (recognizing a fibroblast-specific marker, green) and of α-smooth muscle actin (αSMA: CAF-specific marker, red), comparing DMEM and Mel-MPM. Nuclei are stained blue with Hoechst 33342. E) Vascular tube formation assay of HMEC-1 cells in MCDB131 compared to Mel-MPM in phase contrast and fluorescent mode (merge of Hoechst33342/Calcein AM).

We also tested our melanoma cell lines for their ability to grow as spheroids in ultra-low attachment round bottom plates since we intended to compare 2D and 3D culture systems. Of 24 melanoma cell lines tested, six formed denser spheres in MPM compared to standard cell culture media (Figure S1C and data not shown).

### Establishment of the melanoma-oriented modular physiologic medium (Mel-MPM) for use in more complex, multi-cell-type 3D systems

One aim of the study was the establishment of a matrix-embedded multi-cell-type melanoma 3D model also comprising NHDF fibroblasts, and HMEC-1 endothelial cells. To ensure growth in 3D we supplemented MPM with additional factors. Mel-MPM consists of MPM supplemented with xeno-free B27 supplement (Gibco) and with several growth factors to mimic physiological conditions more closely. The B27 supplement provides additional protein (albumin)-adsorbed nutrients such as essential fatty acids, hormones, lipophilic vitamins, transferrin-bound iron and ROS detoxifying enzymes (catalase, superoxide dismutase) (see Table S2B). The supplemented growth factors were insulin (500 ng/mL), HGF (50 ng/mL), BMP4 (20 ng/mL), EGF (5 ng/mL), and hydrocortisone (1 μg/mL), of which EGF and hydrocortisone are needed for HMEC-1 culture. Those growth factors were selected from a series of preliminary experiments in which we tested various growth factors described to be involved in melanoma tumour growth (data not shown). Importantly, 3D melanoma spheroid growth was increased in Mel-MPM compared to MPM (Figure 1B).

NHDF fibroblasts showed a better 2D growth in Mel-MPM compared to DMEM (Figure 1C). NHDF 3D spheroids shrank in DMEM and showed increased cell death when stained with propidium iodide (PI), while in Mel-MPM the spheroids grew over time and showed little cell death (Figure S1D). NHDF cells in 2D culture displayed no phenotype changes when cultured in Mel-MPM instead of DMEM and presented no changes in immunocytochemical analysis when stained with antibodies for a fibroblast marker (TE7) or alpha smooth muscle actin (αSMA) (Figure 1D). All cells were positive for the TE7 fibroblast marker while only few cells were αSMA-positive.

HMEC-1 vascular cells showed a reduced but still robust 2D growth in Mel-MPM compared to their custom medium, MCDB131 (Figure 1C). Spheroids of HMEC1 showed a slightly better viability in MCDB131 than in Mel-MPM, while the sphere volume was not influenced (Figure 1C and S1D). The ability of HMEC-1 cells to form vascular tubes was identical in MCDB131 and Mel-MPM (Figure 1E). The cells in the vascular tubes were viable as demonstrated by the green live cell stain Calcein AM.

### 2D and 3D cell growth and viability are affected by cystine/cysteine and serine/glycine withdrawal

Taking advantage of the modular composition of MPM, we investigated the consequences of a total withdrawal of individual non-essential amino acids (NEAA, comprising Q and the amino acids present in solution 3: E, N, D, C, CC, S, G), using 2D growth assays (96-well format) with the drug-sensitive 624Mel and WM3248 melanoma cells and derived 624Mel-DR and WM3248-DR cells which are resistant against Encorafenib and Binimetinib (Figure S2). Cystine/cysteine (“MPM-C”) and serine/glycine (“MPM-S/G”) were depleted in combination, while the other amino acids were depleted individually. All four cell lines showed reduced cell numbers upon withdrawal of cystine/cysteine (-C) or serine/glycine (-S/G) while the presence of the other tested NEAA was dispensable for growth (Figure S2). WM3248-DR cells showed a drastically increased cell death in the -C condition and, to a lesser extent, in the -S/G condition. These effects could not be further enhanced by leaving out all non-essential amino acids present in stock solution 3 (-sol3).

We validated the effects (Figure S2) for the cystine/cysteine and serine/glycine starvation conditions in 2D cultures for WM3248 and 624Mel drug-sensitive and drug-resistant cells. Moreover, we included an additional pair of cells: A375 and A375-DR. We grew the cells with different concentrations of C or S/G, keeping the concentration of all other components constant (Figure 2A). The effects observed with the different cell lines were variable. Cystine/Cysteine starvation led to growth reduction in all drug-sensitive and drug-resistant cell lines. Upon cysteine/cystine starvation, WM3248-DR cells show increased Sytox-orange-positive/dead cells in comparison to parental WM3248 cells. On the other hand, 624Mel and 624Mel-DR cells both show moderately decreased cell numbers but no increased Sytox-orange staining. A375 and A375-DR were also very sensitive to cystine/cysteine starvation and showed increased dead cell numbers. Finally, also serine/glycine starvation revealed an increased sensitivity of WM3248-DR and A375-DR cells when compared to their drug-sensitive counterparts with increased numbers of dead cells. In 624Mel and 624Mel-DR cells no such difference in sensitivity was observed upon serine/glycine starvation, the effect of growth suppression was very moderate and dead cell numbers were not upregulated.

**Figure 2.**
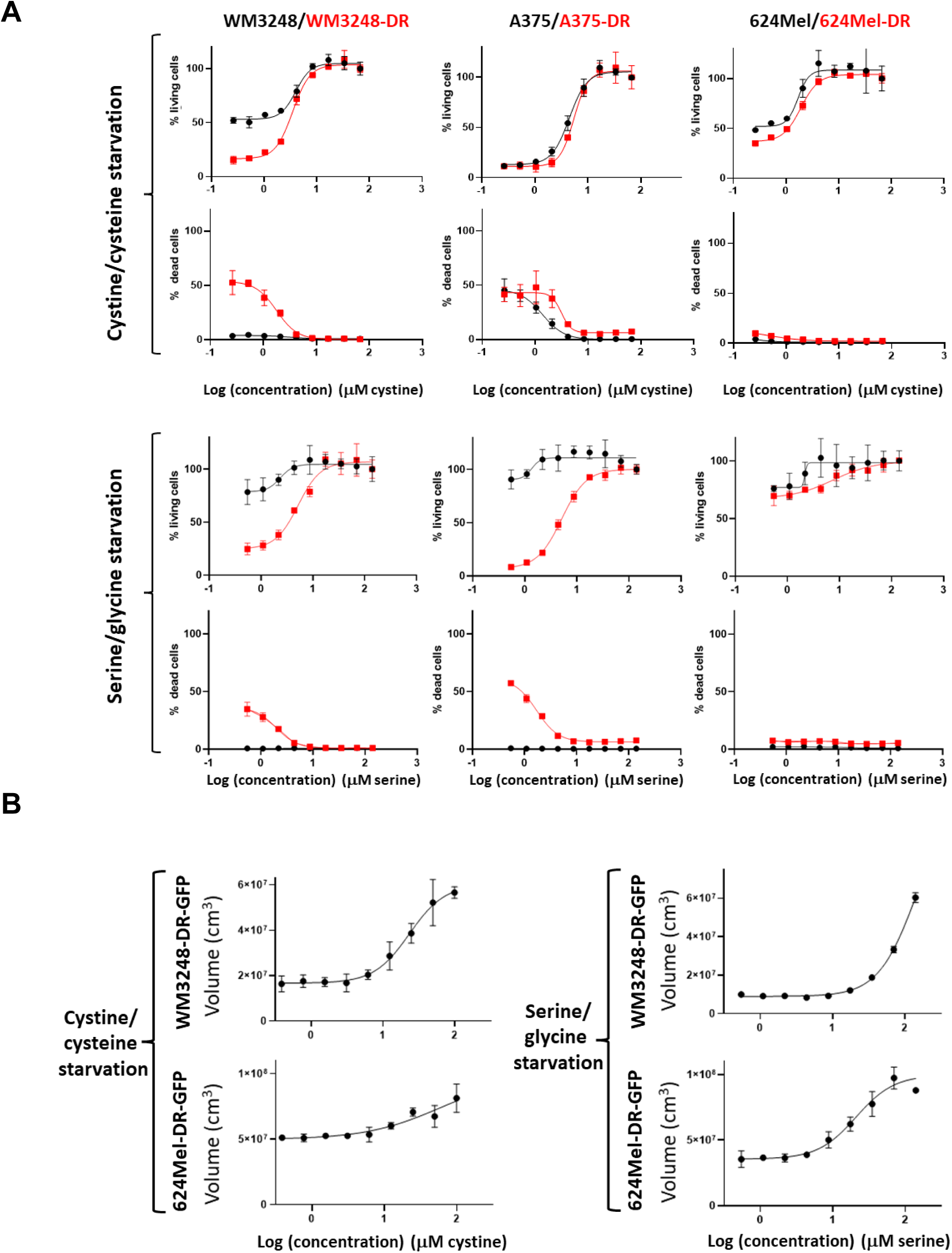
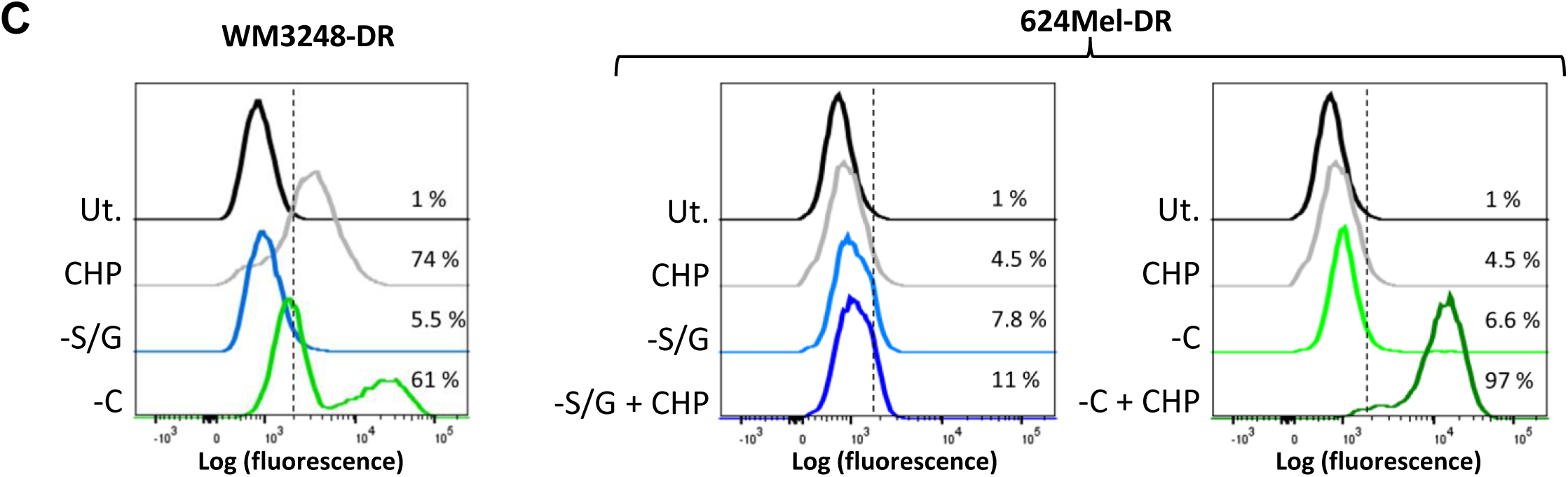
Effect of amino acid starvations on cell viability and ferroptosis induction. A) 2D cultures of the indicated drug-sensitive or drug-resistant melanoma cells were incubated with decreasing concentrations of either cystine/cysteine or serine/glycine (two-fold dilutions starting from the physiological concentrations in MPM; the x-axis for the S/G condition indicates the serine concentration and the x-axis of the cystine/cysteine condition indicates the cystine concentration). Numbers of living or dead cells were determined by using a nuclear cell stain (Hoechst 33342) and a dead cell stain (Sytox orange) using a Cytation 5 instrument and the Gen5 software. B) 3D spheroid cultures of the drug-resistant GFP-expressing WM3248-DR-GFP and 624Mel-DR-GFP cells were treated with decreasing concentrations of either cystine/cysteine or serine/glycine. Spheroid volume was tracked to monitor cell growth. C) Detection of ferroptosis using BODIPY 581/591 C11 for the indicated drug-resistant cells upon 20h of cystine/cysteine (-C) or serine/glycine starvation (-S/G). Cumene hydroperoxide (CHP, 100μM) was used as positive control for ferroptosis induction. Ut.: untreated.

We then investigated 3D spheroid growth in Mel-MPM for the different starving conditions. For these assays, we used GFP-labelled WM3248-DR-GFP and 624Mel-DR-GFP cells. Spheroid volume, GFP expression, and PI uptake by permeabilized dead cells was monitored to investigate the growth and viability of the cells in the spheroids. WM3248-DR-GFP cells showed an efficient growth reduction and cell death induction for the cysteine/cystine and the serine/glycine starvation conditions (Figure 2B). 624Mel-DR-GFP cells showed a less efficient growth reduction and cell death induction for the cystine/cysteine starvation but instead respond better to the serine/glycine starvation treatments compared to 2D cultures.

### Cystine/cysteine starvation alone does not readily induce ferroptosis in MPM

Cystine and glycine are needed to produce glutathione (GSH). Glutathione peroxidase 4 (GPX4) oxidises GSH to GSSG while detoxifying lipid peroxides to the corresponding alcohols, thereby preventing ferroptosis (Figure 3D). In addition, cysteine starvation is known to induce ferroptosis in some cells (Lei et al., 2022). Thus, we investigated whether the increased cell death we observe for cystine/cysteine and serine/glycine starvation in WM3248-DR was due to ferroptosis induction. We determined the levels of the oxidized form of the fluorophore C11-BODIPY, a measure for lipid peroxidation. As a positive control, the ferroptosis inducer cumene hydroperoxide (CHP) was used. Serine/glycine withdrawal did not induce C11-BODIPY oxidation in any cell line. However, in WM3248-DR cells, C11-BODIPY peroxidation was increased by cystine/cysteine withdrawal (Figure 2C, left panel). We did not observe C11-BODIPY oxidation in 624Mel-DR for any single treatment, not even for the positive control (CHP) and we could only induce C11-BODIPY oxidation when cystine/cysteine withdrawal was combined with CHP treatment (Figure 2C, right panels). Thus, although cysteine/cystine withdrawal induces growth reduction in all six and cell death in three of the cell lines tested, C11-BODIPY oxidation, indicative of ferroptosis, is not readily induced by cystine/cysteine withdrawal in MPM medium and is only observed in WM3248-DR cells.

**Figure 3.**
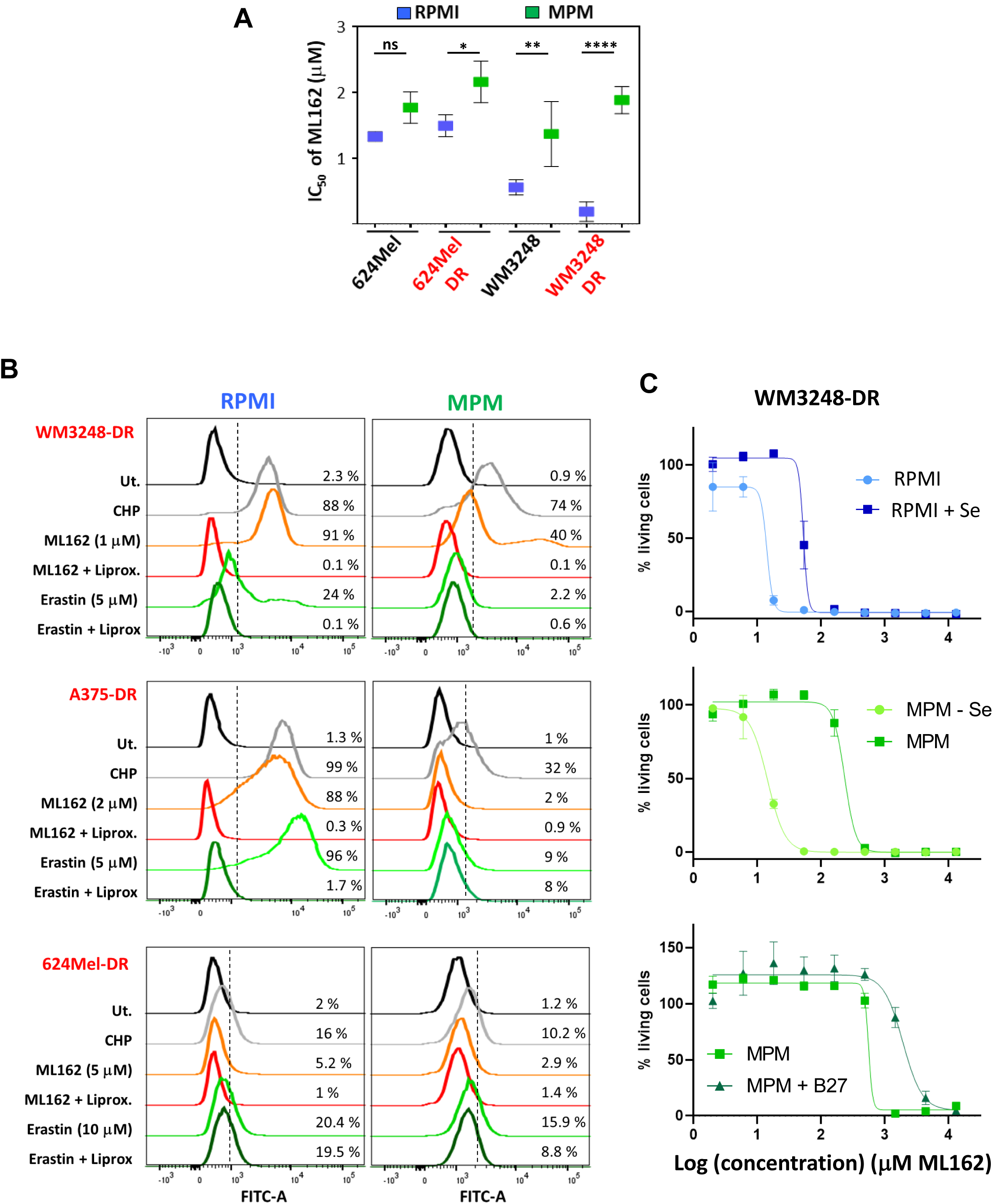
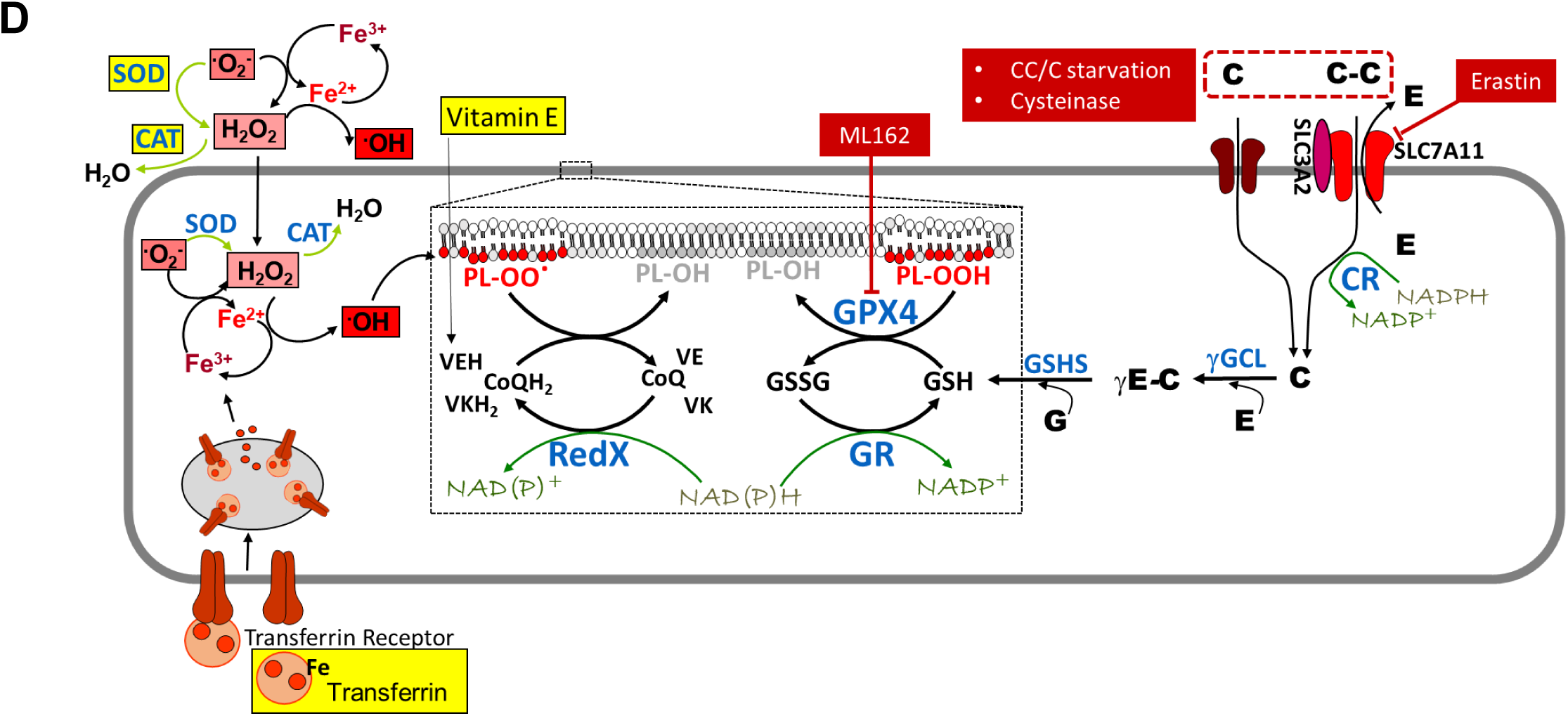
Cell death induction by ML162 can be modulated by the medium composition and by acquired drug resistance. A) The IC50 values of ML162 from CyQuant growth assays performed in RPMI (blue) and in MPM (green) are indicated for 624Mel and WM3248 sensitive and resistant cells. Mean and standard deviation of 3 biological replicates are shown. P values refer to a two-way Anova with Sidaks correction for multiple comparison. NS: P=0.1794, *: P=0.0205, **: P=0.0071, ****: P<0.0001. B) Assessment of ferroptosis induction using BODIPY 581/591 C11 for WM3248-DR, A375-DR, and 624Mel-DR cells upon incubation with ML162 or Erastin (as indicated in the Figure). Cumene hydroperoxide (CHP, 100μM) was used as positive control for ferroptosis induction. Liproxstatin (2 μM) was used as an inhibitor of ferroptosis. C) Growth inhibition of WM3248-DR cells mediated by ML162 is attenuated by the presence of selenite and B27 supplements. IC50 values for ML162 were determined for WM3248-DR cells cultured in different versions of RPMI and MPM supplemented with 10% dialysed FBS. RPMI was either supplemented or not with selenite at the same concentration as used in MPM (0.029 μM). MPM was deprived (or not) of selenite. B27 supplements were added to MPM at the same concentration as used in Mel-MPM, but without adding the growth factors present in Mel-MPM. Prior to the experiment, the WM3248-DR cells were cultured for 3 days in the respective medium. D) The scheme summarises the process of ferroptosis in the light of the different treatments used in this study. (Note: This scheme does not resolve the fact that some enzymes are both present in the cytoplasm and in mitochondria while others are only present in one compartment). Enzymes are indicated in blue. Amino acids (one-letter code) are depicted in fat letters. Inhibitors or starvation treatments are indicated in red-brown boxes with white writing. Components included in B27 are boxed in yellow. Reactive oxygen species are represented by their chemical formula; a more intense red color of boxes indicates higher reactivity. SOD: Superoxide dismutase, CAT: catalase, IDH: isocitrate dehydrogenase, 1C-Met: one carbon metabolism, PPP: pentose phosphate pathway, GPX4: glutathione peroxidase 4, RedX: “reductase X”, indicating different enzymes that can catalyse the reaction (e.g. DHODH in mitochondria, FSP1 in the cytosol), VE: vitamin E, VK: vitamin K, CoQ: coenzyme Q10, PL-OOH and PLOO.. phospholipid peroxide and radical, CR: cystine reductase, SLC7A11: Solute Carrier Family 7 Member 11, SLC3A2: Solute Carrier Family 3 Member 2, γGCL: γ-Glutamate cysteine ligase, GSHS: Glutathione synthetase, NAD: Nicotinamide adenine dinucleotide.

### Cells cultured in physiologic media are more resistant to ferroptosis inducing compounds, such as ML162, compared to cells grown in RPMI

As mentioned above, GPX4 regulates ferroptosis (Lei et al., 2022) and thus, we further tested the effect of the GPX4 inhibitor, ML162, on our cell lines. We determined the IC50 values of ML162 in our melanoma lines in 2D growth assays, comparing RPMI and MPM. In both the 624Mel and WM3248 cells, ML162 was more efficient in suppressing cell numbers in RPMI (Figure 3A). The effect was moderate for 624Mel cells but more pronounced in WM3248 cells. The difference in IC50 values was also more pronounced in drug-resistant cells compared to drug-sensitive cells. Notably, the IC50 values in WM3248 and in WM3248-DR cells in RPMI medium were clearly submicromolar, showing that this ferroptosis-prone cell line was very sensitive, especially in RPMI medium and much less so in MPM.

Next, we measured ferroptosis induction upon ML162 and Erastin treatments in cells cultured in RPMI versus MPM medium. Erastin is an SLC7A11 inhibitor that was also described to induce ferroptosis (Lei et al., 2022). The ferroptosis inducer CHP was again included as a positive control. Liproxstatin, a radical scavenger that prevents lipid peroxidation in membranes, was used to suppress ferroptosis. In WM3248-DR cells cultured in RPMI, C11-BODIPY peroxidation was induced much more efficiently by ML162 than by Erastin (Figure 3B, upper panels) and could in both cases be suppressed by liproxstatin co-treatment. In MPM, however, the C11-BODIPY peroxidation was much less efficient for ML162 than in RPMI, and no signal was observed for Erastin in MPM. For A375-DR cells cultured in RPMI, C11-BODIPY peroxidation was efficiently induced by both ML162 and Erastin (Figure 3B, middle panels) and could in both cases be suppressed by liproxstatin co-treatment. In MPM, however, the C11-BODIPY peroxidation was not observed for any of the two treatments. 624Mel-DR cells proved again to be resilient to ferroptosis induction (Figure 3B, lower panels) and showed no C11-BODIPY oxidation in either RPMI or MPM. Thus, we observe a cell line insensitive to ferroptosis (624Mel-DR), one sensitive in RPMI but not in MPM (A375-DR) and one which shows reduced sensitivity to ML162-induced ferroptosis in MPM (WM3248-DR).

GPX4 is a selenoprotein and it is known that selenite concentrations in media influence protein expression levels of GPX4 (Vande Voorde et al., 2019). Thus, to further investigate the mechanism underlying the different ML162 sensitivity in RPMI *vs*. MPM for the WM3248-DR cells, we pre-cultured those cells in RPMI, RPMI+Se (containing the amount of selenite present in MPM), MPM-Se (MPM without selenite), MPM, and MPM+B27 (the components relevant to ferroptosis in B27 are highlighted in yellow in Figure 3D). After a minimum of 3 days pre-culturing in the different media, we obtained dose-response curves of ML162 in 2D viability assays for WM3248-DR cells. ML162 was suppressing cell viability at lower concentrations in RPMI and MPM when selenite was absent (Figure 3C, upper and middle panel). In addition, ML162 was less efficiently inducing cell death in MPM+B27 (Figure 3C, lower panel).

### ML162 treatment induces apoptosis in cells not undergoing ferroptosis

In preliminary analyses we observed that ML162 treatment could increase caspase 3 activity in 624Mel and WM3248 cells (Figure S3A), which indicated that ML162 induces apoptosis. Investigating caspase-dependent cleavage of PARP by Western blot showed that PARP cleavage occurred in 624Mel, 624Mel-DR and WM3248 cells while it was generally weaker in WM3248-DR, which is the most ferroptosis-prone cell line (Figure S3B).

To further investigate which type of cell death is implicated in ML162-induced effects, we used a microscopic assay detecting the Hoechst 333342 (a live cell fluorescent nuclear stain) and SYTOX orange (a non-cell membrane-permeable fluorescent dead cell dye) fluorescence signals, to track dead cells upon ferroptosis, necrosis or later stage apoptosis. The different cells were also exposed for 16 hours to single treatments or co-treatments with ML162 and various cell death inhibitors: we used Liproxstatin 1, a lipid bilayer-localised antioxidant (LIP: 1μM) and Trolox, a vitamin E analog (TLX: 80 μM), to suppress ferroptosis. Z-VAD-FMK (zVAD: 50 μM) was used to inhibit caspase 3-dependent apoptosis while Necrostatin-1 (NEC: 10 μM), a RIPK1 inhibitor, was used to repress necroptosis. Figure 4 represents the percentage of dead cells for each treatment. Importantly, none of the inhibitors of the different types of cell death leads to important changes on the percentage of dead cells when used alone (Figure 4).

**Figure 4.**
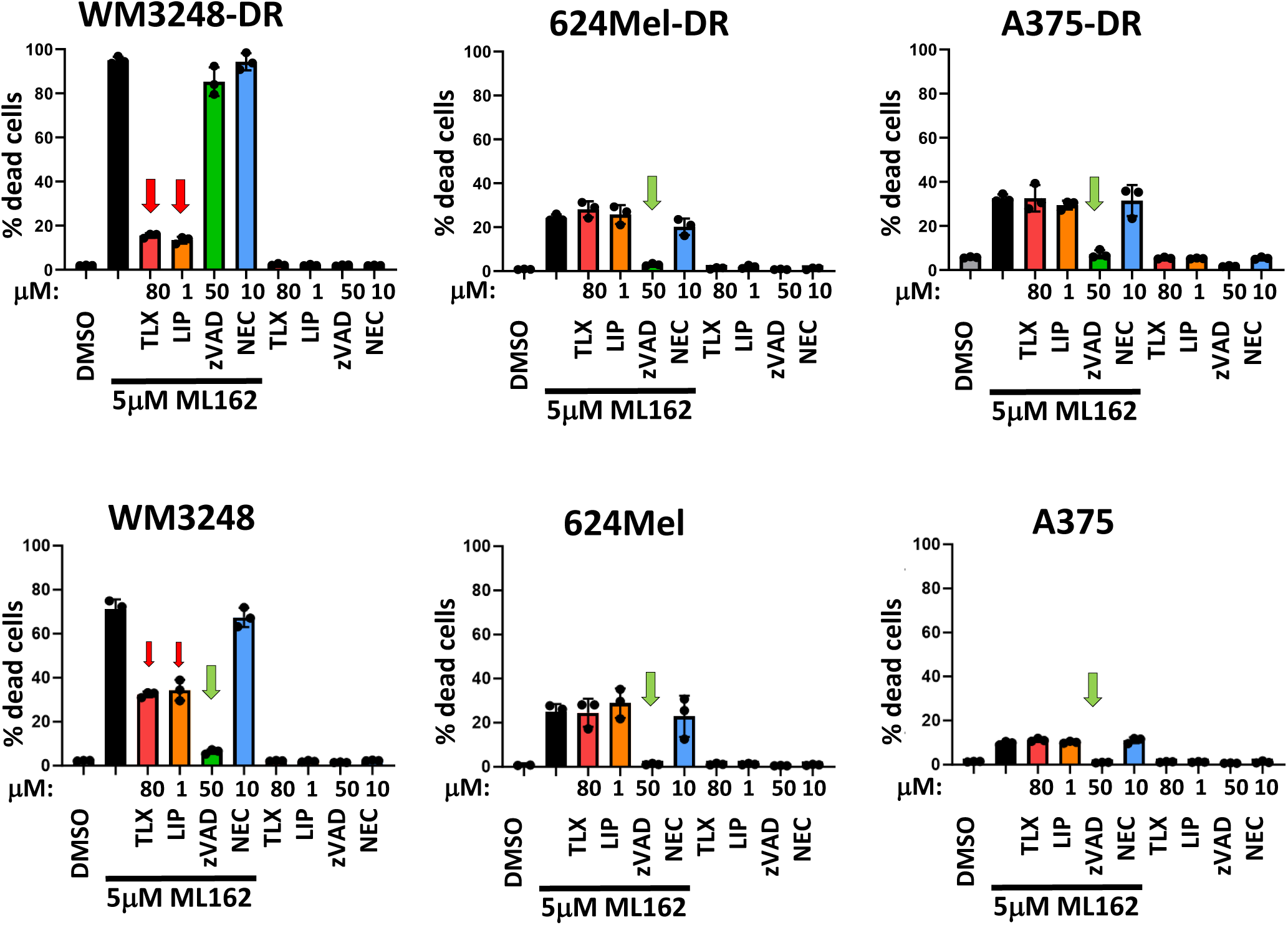
ML162 induces cell death that can be inhibited cell type-dependently by inhibitors of ferroptosis or apoptosis. 2D cell cultures were grown in MPM with 5μM ML162 and with various inhibitors of different cell death inducing pathways, as follows: LIP: Liproxstatin 1 (1μM), TLX: Trolox (80μM), zVAD: Z-VAD-FMK (50μM), NEC: Necrostatin-1 (10μM). The numbers of viable and dead cells were determined microscopically with the Cytation 5 instrument using Hoechst 33342 to stain the nuclei of live and dead cells and using Sytox orange to stain dead cells. Red arrows indicate the ferroptosis inhibitor (Liproxstatin 1, Trolox) effects. Green arrows indicate the caspase (apoptosis; Z-VAD-FMK) inhibitor effect.

In MPM, WM3248-DR cells show a very prominent increase in dead cells when treated with ML162, which was rescued by cotreatment with the two ferroptosis inhibitors (left upper graph, red arrows), while the apoptosis and necroptosis inhibitors did not reduce the percentage of dead cells. Interestingly, WM3248 show a partial rescue by the ferroptosis inhibitor (left lower graph, thin red arrows) and by the apoptosis inhibitor (left lower graph, green arrow), while the apoptosis inhibitor (green bar) seems to play the major role in rescuing WM3248 cells. All other cells, A375, A375-DR, 624Mel and 624Mel-DR, showed a suppression of dead cells when ML162 was co-administered with the caspase inhibitor zVAD (see green arrows). It should be noted that the use of zVAD only inhibits caspase 3 activity, which ultimately also leads to secondary necrosis but does not counteract the initial toxic effect of ML162. This may explain why the number of live cells is still reduced in this situation (data not shown). The necroptosis inhibitor showed no rescuing effects in any cell line. Thus, ML162-induced apoptosis seems to play a major role in 5 of our cell lines, while only one cell line, WM3248-DR, undergoes ferroptosis in MPM.

### ML162 treatment induces changes in mitochondrial ROS

Western blot analysis of 624Mel and 624Mel-DR cells also showed that 2 hours after treatment with ML162, the NRF2 protein is upregulated, which is induced after oxidative stress (Figure S3C). ML162 treatment might thus lead to mitochondrial stress by increasing ROS, which might induce apoptosis via the intrinsic pathway of apoptosis (Figure 5A). Indeed, upon stimulation of cells with ML162, an increase in the MitoSox-green stain was observed, indicating an increase in mitochondrial superoxide while the overall mitochondrial mass did not change (detected using MitoLite red) (Figure 5B).

**Figure 5.**
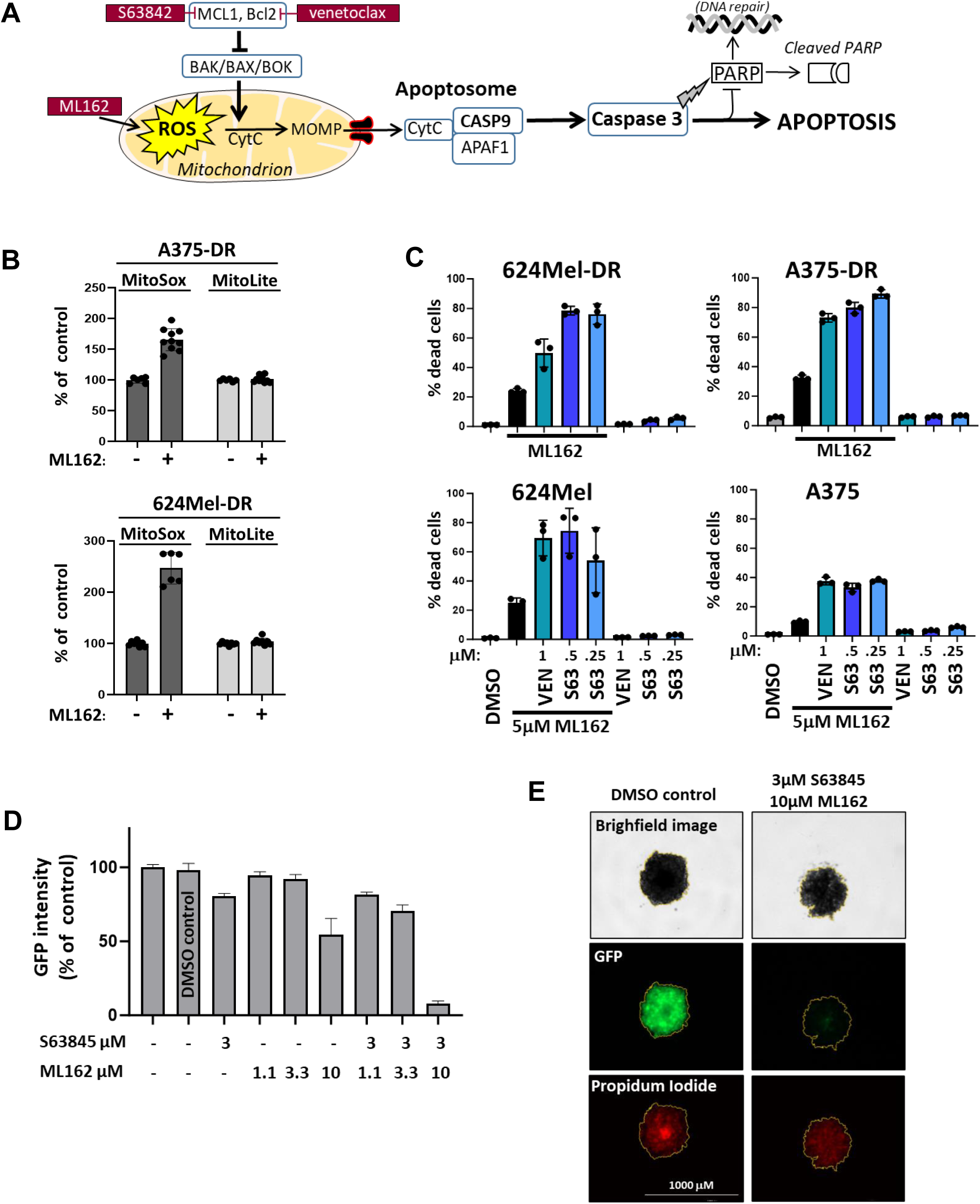
Co-treatment of ML162 with compounds triggering the intrinsic apoptosis pathway leads to increased cell death. A) Scheme showing the intrinsic apoptosis pathway and how it could be triggered by ML162 and other compounds. B) A375-DR and 624Mel-DR cells were left untreated or were treated with 5 μM ML162 for 50 minutes. Mitochondrial superoxide production was measured using MitoSox-Green (5 uM for A375-DR, 7,5 uM for 624Mel-DR). MitoLite-Red (diluted to 0.8X) was used as control to see possible changes in overall mitochondrial mass. C) Cells grown in 2D were treated with 5 μM ML162 and with inhibitors of BCL2 (Venetoclax: VEN) and MCL1 (S63845: S63) to induce apoptosis via the intrinsic pathway. The numbers of viable and dead cells were determined microscopically with the Cytation 5 instrument using Hoechst 33342 to stain the nuclei of live and dead cells and Sytox orange to stain dead cells. D) 3D spheroids of 624Mel-DR-GFP cells were exposed to the MCL1 inhibitor S63845 and ML162, as indicated. To assess the viability of the cells in 3D spheroids, the GFP expression in the cells was measured. E) Representative pictures of some conditions of the 3D spheroid assay described in D).

### Cotreatment with Bcl2 and MCL1 inhibitors efficiently increases the effect of ML162 treatment on the induction of apoptosis in 2D and 3D assays

Since ML162 seems to induce ROS in mitochondria, it might sensitize cells to apoptosis *via* the intrinsic pathway of apoptosis (Figure 5A). We tested whether inhibitors of BH1-4 proteins, such as the BCL2 inhibitor Venetoclax, or the MCL1 inhibitor S63845, can increase the effect of ML162 on our drug-sensitive and -resistant melanoma cells. At concentrations where these compounds alone had no effect on cell viability, they reduced survival of the melanoma cell lines when used together with ML162 (Figure 5C). The MCL1 inhibitor especially had good potency and could be used at submicromolar levels in 2D assays. We then determined how this combination works in 3D spheroid assays of 624Mel-DR cells. Preliminary tests showed that the concentrations of both drugs had to be raised to have effects in 3D (data not shown). Figure 5D and 5E show the effect of combinations of 3 μM S63845 with varying concentration of ML162 on 624Mel-DR-GFP 3D spheroids. The combination of both increases the effect of ML162 on cell viability considerably (measured here by following GFP expression), but both drugs have to be used at higher concentrations compared to the 2D assays. Since the concentrations could not be lowered below the threshold at which ML162 shows toxic effects in fibroblasts and endothelial cells (see last results paragraphs), we conclude that this combination does very likely not represent an efficient treatment strategy for melanoma and do not discuss this further.

### Cystine/cysteine starvation sensitizes ferroptosis-prone WM3248-DR cells to ML162 treatment but fails to do so in A375-DR and 624Mel-DR cells

To test if additional cystine/cysteine starvation sensitizes drug-resistant melanoma cells to the GPX4 inhibitor ML162, we incubated the drug-resistant lines with a cystine/cysteine concentration at which growth was already affected but cells still retained a viability of around 70 to 90% (see Figure 6A for the exact concentrations). Then, ML162 was added at different concentrations for 2D viability assays. Figure 6A shows that the ferroptosis-prone WM3248-DR cells were sensitized to ML162 treatment when grown in MPM containing only 10% of the physiological cystine/cysteine concentration in human plasma (6.7 μM cystine and 3.2 μM cysteine). However, the apoptosis-prone 624Mel-DR and A375-DR cells could not be sensitized to ML162 treatment.

**Figure 6.**
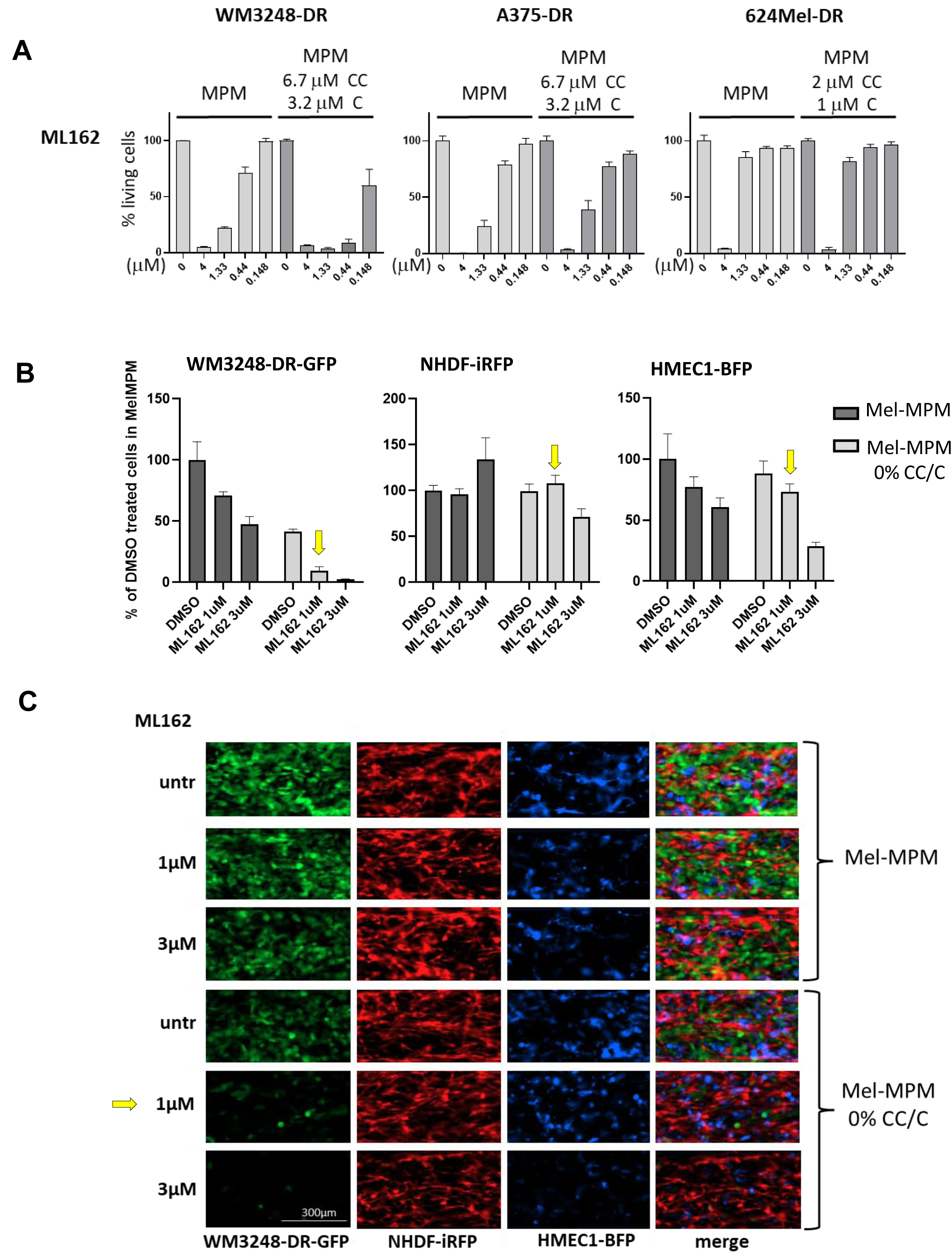
Effect of co-treatments (ML162 treatment and cystine/cysteine reduction) on drug-resistant melanoma cells. A) Drug-resistant melanoma cells, grown in 2D, were incubated in either MPM or MPM containing 6.7 μM cystine and 3.2 μM cysteine. ML162 was added at the indicated concentrations using half-log dilution steps. Live cell numbers were determined using a nuclear cell stain (Hoechst 33342) and a dead cell stain (Sytox orange). Images were acquired on a Cytation 5 instrument and evaluated using the Gen5 software. Bars represent the percentage of living cells relative to the respective non-drug-treated condition. B) and C) Quantitation of the different fluorescent signals (B) and images (C) of matrix-embedded co-cultures of WM3248-DR cells with NHDFs and HMEC-1 cells in OrganoPlates™. 10,000 WM3248-DR-GFP, 5,000 NHDF-iRFP, and 5,000 HMEC1-BFP per μL Geltrex were seeded in the OrganoPlate. After 2 days, the medium was replaced, and cells were either left untreated or treated with 1 or 3 μM ML162 or with a combination of cysteine deprivation (0% CC and C) and ML162. 4 days later, the cells were imaged on a Cytation 10 instrument using a 10x objective and the LED/Filter cubes for GFP, Cy5.5, and DAPI. The yellow arrow highlights the condition in which melanoma cell numbers are suppressed specifically compared to NHDF and HMEC1.

### Combination of cystine/cysteine withdrawal and ML162 treatment in a matrix-embedded multi-cell-type 3D system shows that drug concentrations can be optimized to preferentially target the growth of ferroptosis-prone melanoma cells

We then tested if the cystine/cysteine starvation and ML162 treatment can be combined to potently suppress melanoma growth in ferroptosis-prone WM3248-DR cells in a matrix-embedded multi-cell-type melanoma model. For this purpose, we used GFP-labelled WM3248-DR cells, iRFP-labelled neonatal human dermal fibroblasts (NHDF-iRFP) and tagBFP-labelled human microvessal endothelial cells (HMEC1-BFP) (Figure 6B). WM3248-DR-GFP, HMEC1-BFP, and NHDF-iRFP cells were embedded in Geltrex matrix and seeded into the microchannels of an OrganoPlate™ (Mimetas; for a detailed representation of cell growth in the matrix-containing microchannel, see Figure S4). The different cell types can be visualized according to their specific fluorophores (expressed in viable cells) to assess the effects of the drug treatments on the different cell types. After 2 days of pre-incubation, the medium was replaced with either Mel-MPM or starvation medium (Mel-MPM without cystine/cysteine) which did or did not contain drugs. After 4 days of treatment with1 μM ML162, the viability of the WM3248-DR cells was drastically reduced when combined with cystine/cysteine withdrawal, while this condition had no or moderate effects on NHDF-iRFP and HMEC1-BFP viability (see marked condition in Figures 6B/C). This observation suggests that, while physiologic media generally reduce the sensitivity of drug-resistant cells for ferroptosis inducing stimuli, some ferroptosis-prone cell lines, such as WM3248, can still be targeted to undergo ferroptosis at conditions that leave other cell types unscathed.

## Discussion

Many compounds identified in preclinical *in vitro* models fail to enter clinical trials, since classical *in vitro* models (cell lines grown as monolayer in standard cell culture media) do not reflect important features of tumours, which are characterized by a complex distribution of different metabolic and cellular environments, resulting in complex patterns of proliferative, quiescent, and necrotic regions (Sant & Johnston, 2017). There is a need to develop more physiological cell culture systems suitable for screening approaches. These would optimally reflect i) the blood, lymph or tissue metabolomes in which cancer cells thrive (physiologic media), ii) the heterogeneity of tumours represented by oxygen and nutrient gradients (3D cell culture and/or oxygen control), (iii) the cellular complexity of tumours by containing cell types of other tissue origin (e.g. fibroblasts, endothelial cells), and iv the matrix composition and stiffness in which tumour cells grow. Consequently, we aimed to develop a physiologic medium, adaptable in composition, for melanoma 2D and 3D cell growth, and to further establish models which are “screenable” for nutrient dependencies and drug sensitivities, using also matrix-embedded multi-cell-type 3D cultures (see Figure S4).

There is increasing evidence that metabolic processes are different in physiologic versus standard media, as reviewed recently (Cantor, 2019; Golikov et al., 2022). A comparison of metabolic profiles from a large numbers of cell lines cultured either in DMEM or RPMI established that over 60% of metabolites measured were correlated significantly with the medium type used (Li et al., 2019;Lagziel et al., 2020). In line with this, CRISPR- or RNA-interference-gene silencing approaches or compound screens were described to yield different results when performed in different media, especially when targeting metabolic enzymes (Abbott et al., 2023; Lagziel et al., 2019;Rossiter et al., 2021). Therefore, we implemented a novel physiologic medium (MPM – modular physiologic medium), better representing the nutrient concentrations in plasma. MPM is similar to the formulations of HPLM and Plasmax (Cantor et al., 2017; Vande Voorde et al., 2019), although individual metabolite concentrations differ (see Tables S1 and S2). The three mentioned physiologic media contain between 27 to 31 additional metabolites absent in standard media, which make up 35.5 to 50% of the total nutrient concentrations: 35.5% for Plasmax (Vande Voorde et al., 2019), 50% for HPLM (Cantor et al., 2017), and 45% for MPM (Figure S1). The amino acid concentrations in all three physiologic media are slightly different but designed to represent the human plasma amino acid concentrations. In standard cell culture media, nutrients (glucose and amino acids) are mostly present at above physiological concentrations (Ackermann & Tardito, 2019) and greatly vary between the different media (see MEM, RPMI, DMEM, IMDM in Table S1 and Figure S1). For 3D spheroid growth and for the more complex multi-cell-type matrix-embedded culture in Organoplates^TM^, we supplemented MPM with additional factors (Mel-MPM contains xeno-free B27 and additional growth factors).

Taking advantage of the modular build-up of MPM, we tested for vulnerabilities of drug-resistant melanoma cells upon withdrawal of specific non-essential amino acids from MPM. We describe that cystine/cysteine and serine/glycine starvation most efficiently suppressed 2D melanoma cell growth, while cells were resilient towards the single deprivation of other tested NEAA like glutamine, glutamate, asparagine, or aspartate. Cystine/cysteine depletion is known to lower the levels of GSH and to be able to induce ferroptosis (Daher et al., 2020; Jiang et al., 2021;Meinert et al., 2024). In MPM lacking cystine/cysteine, we could induce C11-BODIPY peroxidation in WM3248-DR cells, while it was not possible in 624Mel-DR. Cancers cells are known to rely on extracellular cystine to meet their need for the antioxidant GSH, especially because the *de novo* synthesis of cysteine *via* the transsulfuration pathway or regeneration of cysteine by protein catabolism have been shown to provide insufficient amounts (Chio & Tuveson, 2017; Koppula et al., 2021; Trachootham et al., 2009). In our cells, serine/glycine withdrawal (both amino acids being also involved in GSH production in melanoma (Meinert et al., 2024) did not induce C11-BODIPY peroxidation in MPM (see Figure 2C).

We further used ferroptosis inducing compounds such as the GPX4 inhibitor ML162 and the SLC7A11 inhibitor Erastin to test their cell death inducing capacity in our physiologic medium (Figure 3). For ML162, the IC50 values determined in growth assays in RPMI compared to MPM were much lower for the WM3248 and WM3248-DR cell lines, while they were quite similar in 624Mel and 624Mel-DR. WM3248-DR cells cultured in RPMI showed higher levels of C11-BODIPY peroxidation upon ML162 treatment compared to cells cultured in MPM (Figure 3B, upper panels). For A375-DR cells in which C11-BODIPY peroxidation was efficiently induced by both ML162 and Erastin when cultured in RPMI (Figure 3B, middle panels), it was totally suppressed in MPM. The ML162 target GPX4 is a selenoprotein, and it is known that selenite concentrations in media influence protein expression levels of GPX4 (Vande Voorde et al., 2019). The selenite addition / removal to RPMI and MPM, respectively, showed that ML162 was less effective when selenite was present in the media (Figure 3C). The addition of xeno-free B27, which contains (albumin)-adsorbed nutrients such as essential lipids, hormones, lipophilic vitamins (Vit A and E), transferrin-bound iron, and ROS-detoxifying enzymes (catalase, superoxide dismutase) (Brewer et al., 1993) also led to a reduction in ML162 potency in dose response assays with WM3248-DR cells (Figure 3C). This indicates that ferroptosis inducing compounds are more effective in standard cell media devoid of selenite (such as RPMI or DMEM) compared to more physiologic media that contain selenite, vitamin E, and ROS-detoxifying enzymes (catalase, superoxide dismutase). Thus, we conclude that induction of lipid peroxidation, a hallmark of ferroptosis, by either cystine starvation or ferroptosis inducing compounds, is harder to achieve in the more physiological medium MPM than in RPMI.

We further show that, although physiologic media reduce the effectiveness of ML162 or cystine/cysteine starvation, ferroptosis-prone cell lines can still undergo ferroptosis if multiple treatments are combined. Cell viability of WM3248-DR cells was efficiently and potently reduced when ML162 (1 μM) treatment was combined with cystine/cysteine starvation in 2D (Figure 6A) and in our more complex matrix-embedded multi-cell-type model (Figure 6B/C, Figure S4).

Interestingly, systemic cystine/cysteine depletion has been described to be achievable *in vivo* in mouse models by treatment with cyst(e)inase (Cramer et al., 2017). Generally, this approach could be further optimised by targeting cysteinase specifically to tumours using e.g. cyst(e)inase-linked tumour antigen-targeted antibodies and to combine this with ML162 or other ferroptosis inducing drugs. Thus, cystine/cysteine depletion and drug co-treatment strategies might be a treatment option for cancer in the future, as more and more specific diets for cancer treatment are proposed (Butler et al., 2021; Kanarek et al., 2020; Lien & Vander Heiden, 2019; Muir et al., 2018b; Sullivan & Vander Heiden, 2019; Upadhyayula et al., 2023). This is especially important in the light of data which showed that melanoma cells were more resistant to ferroptosis in lymph compared to blood, which has been attributed to higher concentrations of GSH, oleic acid, and to lower free iron levels in lymph (Ubellacker et al., 2020). Thus, different physiological compartments in which cancer cells thrive, such as blood, lymph or interstitial fluid inside a tumor, present different nutrient compositions and may influence the cancer cells response to therapy. Co-treatment strategies might be especially important to be able to target cancer cells in different physiological niches, in which they might be more protected from single treatment.

Taken together, we have established MPM and Mel-MPM as novel resources to cultivate melanoma cells under conditions “closer to physiology”. We find that ferroptosis is less likely to occur in physiologic medium mimicking blood plasma metabolite concentrations. However, cells can be sensitised to undergo ferroptosis upon drug treatment in cystine/cysteine-depleted conditions. We provide evidence that the novel Mel-MPM medium sustains 3D growth in multi-cell-type matrix-embedded cultures in Organoplates^TM^ containing melanoma cells, NHDFs, and HMEC-1 cells. We use this complex system to show that there is a window of opportunity to treat ferroptosis-prone drug-resistant melanoma cells with cysteine starvation and ML162, without impairing growth of other cell types (Figure 6B/C, Figure S4). In its 96-well format, the Mel-MPM/Organoplates^TM^ co-culture system can be implemented for screening approaches. Our results also underline the power of combining metabolism-oriented drug treatments with amino acid starvation conditions, which is of interest in view of future therapeutic approaches to combat melanoma and other cancer types.

## Acknowledgements

The 624Mel and M45 cells were a kind gift of Dr. Ruth Halaban (Dermatology Department, Yale School of Medicine, USA) and Dr. Dagmar Kulms (Dresden University, Germany), respectively. We thank Dr. Martin Nurmik (University of Luxembourg) for help with the sorting of the GFP expressing melanoma lines. We thank Dr. Heike Hermanns and the Zentrallabor (University Clinic Würzburg, Germany) for Osmolality measurements. This work was supported by the Luxembourg National Research Fund (FNR), project number 10675146 (funding scheme PRIDE, “CANBIO”), and by the Fondation Cancer Luxembourg (“SecMelPro” grant).

## Abbreviations

αSMA: alpha smooth muscle actin

AA: amino acid

BME: Basal Medium Eagle

BMP4: Bone Morphogenetic Protein-4

BFP: blue fluorescent protein

CoQH2: reduced coenzyme Q

CHP: cumene hydroperoxide

DR: drug-resistant

EBSS: Earle’s Balanced Salt Solution

EGF: epidermal growth factor

FBS: fetal bovine serum

eIF2α: eukaryotic initiation factor 2α

FACS: fluorescence-activated cell sorting

FITC: fluorescein isothiocyanate

GAPDH: glyceraldehyde 3-phosphate dehydrogenase

GFP: green fluorescent protein

GSH: glutathione

GSSG: Glutathione disulfide

GPX4: Glutathione peroxidase 4

HGF: Hepatocyte Growth Factor

HPLM: Human Plasma-Like Medium

iRFP: near-infrared fluorescent protein

Mel-MPM: melanoma-oriented modular physiologic medium

MPM: modular physiologic medium

NAD: nicotinamide adenine dinucleotide

NAMPT: nicotinamide phosphoribosyltransferase

NAMPTi: nicotinamide phosphoribosyltransferase inhibitor

NEAA: non-essential amino acid

NHDF: Normal Human Dermal Fibroblasts

PI: propidium iodide

ROS: reactive oxygen species

Se: selenite

SLC3A2: Solute Carrier Family 3 Member 2

SLC7A11: Solute Carrier Family 7 Member 11

xCT: cystine/glutamate transporter

## References

Ackermann, T., & Tardito, S. (2019). Cell Culture Medium Formulation and Its Implications in Cancer Metabolism. Trends in Cancer, 5(6), 329–332. 10.1016/j.trecan.2019.05.004

Alkaraki, A., McArthur, G. A., Sheppard, K. E., & Smith, L. K. (2021a). Metabolic plasticity in melanoma progression and response to oncogene targeted therapies. Cancers, 13(22). 10.3390/cancers13225810

Avagliano, A., Fiume, G., Pelagalli, A., Sanità, G., Ruocco, M. R., Montagnani, S., & Arcucci, A. (2020a). Metabolic Plasticity of Melanoma Cells and Their Crosstalk With Tumor Microenvironment. Frontiers in Oncology, 10(May), 1–21. 10.3389/fonc.2020.00722

Avellino, G., Deshmukh, R., Rogers, S. N., Charnock-Jones, D. S., Smith, G. C. S., Tardito, S., & Aye, I. L. M. H. (2023). Physiologically relevant culture medium Plasmax improves human placental trophoblast stem cell function. American Journal of Physiology - Cell Physiology, 324(4), C878–C885. 10.1152/ajpcell.00581.2022

Brewer, G. J., & Cotman, C. W. (1989). Survival and growth of hippocampal neurons in defined medium at low density: advantages of a sandwich culture technique or low oxygen. Brain Research, 494(1), 65–74. 10.1016/0006-8993(89)90144-3

Brewer, G. J., Torricelli, J. R., Evege, E. K., & Price, P. J. (1993). Optimized survival of hippocampal neurons in B27-supplemented neurobasal^TM^, a new serum-free medium combination. Journal of Neuroscience Research, 35(5), 567–576. 10.1002/jnr.490350513

Butler, M., van der Meer, L. T., & van Leeuwen, F. N. (2021). Amino Acid Depletion Therapies: Starving Cancer Cells to Death. Trends in Endocrinology and Metabolism, 32(6), 367–381. 10.1016/j.tem.2021.03.003

Cantor, J. R. (2019). The Rise of Physiologic Media. Trends in Cell Biology, 29(11), 854–861. 10.1016/j.tcb.2019.08.009

Cantor, J. R., Abu-Remaileh, M., Kanarek, N., Freinkman, E., Gao, X., Louissaint, A., Lewis, C. A., & Sabatini, D. M. (2017). Physiologic Medium Rewires Cellular Metabolism and Reveals Uric Acid as an Endogenous Inhibitor of UMP Synthase. Cell, 169(2), 258–272.e17. 10.1016/j.cell.2017.03.023

Chio, I. I. C., & Tuveson, D. A. (2017). ROS in Cancer: The Burning Question. Trends in Molecular Medicine, 23(5), 411–429. 10.1016/j.molmed.2017.03.004

Cramer, S. L., Saha, A., Liu, J., Tadi, S., Tiziani, S., Yan, W., Triplett, K., Lamb, C., Alters, S. E., Rowlinson, S., Zhang, Y. J., Keating, M. J., Huang, P., DiGiovanni, J., Georgiou, G., & Stone, E. (2017). Systemic depletion of L-cyst(e)ine with cyst(e)inase increases reactive oxygen species and suppresses tumor growth. Nature Medicine, 23(1), 120–127. 10.1038/nm.4232

Daher, B., Vučetić, M., & Pouysségur, J. (2020). Cysteine Depletion, a Key Action to Challenge Cancer Cells to Ferroptotic Cell Death. Frontiers in Oncology, 10(May), 1–9. 10.3389/fonc.2020.00723

Dereziński, P., Klupczynska, A., Sawicki, W., Pałka, J. A., & Kokot, Z. J. (2017). Amino acid profiles of serum and urine in search for prostate cancer biomarkers: A pilot study. International Journal of Medical Sciences, 14(1), 1–12. 10.7150/ijms.15783

Dummer, R., Flaherty, K. T., Robert, C., Arance, A., De Groot, J. W. B., Garbe, C., Gogas, H. J., Gutzmer, R., Krajsová, I., Liszkay, G., Loquai, C., Mandalà, M., Schadendorf, D., Yamazaki, N., Di Pietro, A., Cantey-Kiser, J., Edwards, M., & Ascierto, P. A. (2022). COLUMBUS 5-Year Update: A Randomized, Open-Label, Phase III Trial of Encorafenib Plus Binimetinib Versus Vemurafenib or Encorafenib in Patients With BRAF V600-Mutant Melanoma. Journal of Clinical Oncology, 40(36). 10.1200/JCO.21.02659

Golikov, M. V., Valuev-Elliston, V. T., Smirnova, O. A., & Ivanov, A. V. (2022). Physiological Media in Studies of Cell Metabolism. Molecular Biology, 56(5), 629–637. 10.1134/S0026893322050077

Haan, C., & Behrmann, I. (2007). A cost effective non-commercial ECL-solution for Western blot detections yielding strong signals and low background. Journal of Immunological Methods, 318(1–2), 11–19. 10.1016/j.jim.2006.07.027

Jiang, X., Stockwell, B. R., & Conrad, M. (2021). Ferroptosis: mechanisms, biology and role in disease. Nature Reviews Molecular Cell Biology, 22(4), 266–282. 10.1038/s41580-020-00324-8

Kanarek, N., Petrova, B., & Sabatini, D. M. (2020). Dietary modifications for enhanced cancer therapy. Nature, 579(7800), 507–517. 10.1038/s41586-020-2124-0

Koppula, P., Zhuang, L., & Gan, B. (2021). Cystine transporter SLC7A11/xCT in cancer: ferroptosis, nutrient dependency, and cancer therapy. Protein and Cell, 12(8), 599–620. 10.1007/s13238-020-00789-5

Kozar, I., Margue, C., Rothengatter, S., Haan, C., & Kreis, S. (2019). Many ways to resistance: How melanoma cells evade targeted therapies. BBA - Reviews on Cancer, 1871(2), 313–322. 10.1016/j.bbcan.2019.02.002

Kramer, B., De Haan, L., Vermeer, M., Olivier, T., Hankemeier, T., Vulto, P., Joore, J., & Lanz, H. L. (2019). Interstitial flow recapitulates gemcitabine chemoresistance in a 3D microfluidic pancreatic ductal adenocarcinoma model by induction of multidrug resistance proteins. International Journal of Molecular Sciences, 20(18). 10.3390/ijms20184647

Lagziel, S., Gottlieb, E., & Shlomi, T. (2020). Mind your media. In Nature Metabolism (Vol. 2, Issue 12). 10.1038/s42255-020-00299-y

Lagziel, S., Lee, W. D., & Shlomi, T. (2019). Inferring cancer dependencies on metabolic genes from large-scale genetic screens. BMC Biology, 17(1). 10.1186/s12915-019-0654-4

Lee, S. H., Hu, W., Matulay, J. T., Silva, M. V., Owczarek, T. B., Kim, K., Chua, C. W., Barlow, L. M. J., Kandoth, C., Williams, A. B., Bergren, S. K., Pietzak, E. J., Anderson, C. B., Benson, M. C., Coleman, J. A., Taylor, B. S., Abate-Shen, C., McKiernan, J. M., Al-Ahmadie, H., … Shen, M. M. (2018). Tumor Evolution and Drug Response in Patient-Derived Organoid Models of Bladder Cancer. Cell, 173(2), 515–528.e17. 10.1016/j.cell.2018.03.017

Lei, G., Zhuang, L., & Gan, B. (2022). Targeting ferroptosis as a vulnerability in cancer. Nature Reviews Cancer, 0123456789. 10.1038/s41568-022-00459-0

Li, H., Ning, S., Ghandi, M., Kryukov, G. V., Gopal, S., Deik, A., Souza, A., Pierce, K., Keskula, P., Hernandez, D., Ann, J., Shkoza, D., Apfel, V., Zou, Y., Vazquez, F., Barretina, J., Pagliarini, R. A., Galli, G. G., Root, D. E., … Sellers, W. R. (2019). The landscape of cancer cell line metabolism. Nature Medicine, 25(5). 10.1038/s41591-019-0404-8

Lien, E. C., & Vander Heiden, M. G. (2019). A framework for examining how diet impacts tumour metabolism. Nature Reviews Cancer, 19(11), 651–661. 10.1038/s41568-019-0198-5

Meinert, M., Jessen, C., Hufnagel, A., Kreß, J. K. C., Burnworth, M., Däubler, T., Gallasch, T., Xavier da Silva, T. N., dos Santos, A. F., Ade, C. P., Schmitz, W., Kneitz, S., Friedmann Angeli, J. P., & Meierjohann, S. (2024). Thiol starvation triggers melanoma state switching in an ATF4 and NRF2-dependent manner. Redox Biology, 70. 10.1016/j.redox.2023.103011

Muir, A., Danai, L. V., & Vander Heiden, M. G. (2018a). Microenvironmental regulation of cancer cell metabolism: Implications for experimental design and translational studies. DMM Disease Models and Mechanisms, 11(8). 10.1242/dmm.035758

Pendleton, K. E., Wang, K., & Echeverria, G. V. (2023). Rewiring of mitochondrial metabolism in therapy-resistant cancers: permanent and plastic adaptations. In Frontiers in Cell and Developmental Biology (Vol. 11). 10.3389/fcell.2023.1254313

Psychogios, N., Hau, D. D., Peng, J., Guo, A. C., Mandal, R., Bouatra, S., Sinelnikov, I., Krishnamurthy, R., Eisner, R., Gautam, B., Young, N., Xia, J., Knox, C., Dong, E., Huang, P., Hollander, Z., Pedersen, T. L., Smith, S. R., Bamforth, F., … Wishart, D. S. (2011). The human serum metabolome. PLoS ONE, 6(2). 10.1371/journal.pone.0016957

Rossiter, N. J., Huggler, K. S., Adelmann, C. H., Keys, H. R., Soens, R. W., Sabatini, D. M., & Cantor, J. R. (2021a). CRISPR screens in physiologic medium reveal conditionally essential genes in human cells. Cell Metabolism, 2020.08.31.275107. 10.1016/j.cmet.2021.02.005

Sant, S., & Johnston, P. A. (2017). The production of 3D tumor spheroids for cancer drug discovery. Drug Discovery Today: Technologies, 23(xx), 27–36. 10.1016/j.ddtec.2017.03.002

Sullivan, M. R., & Vander Heiden, M. G. (2019). Determinants of nutrient limitation in cancer. Critical Reviews in Biochemistry and Molecular Biology, 54(3), 193–207. 10.1080/10409238.2019.1611733

Torres-Quesada, O., Doerrier, C., Strich, S., Gnaiger, E., & Stefan, E. (2022). Physiological Cell Culture Media Tune Mitochondrial Bioenergetics and Drug Sensitivity in Cancer Cell Models. Cancers, 14(16). 10.3390/cancers14163917

Trabado, S., Al-Salameh, A., Croixmarie, V., Masson, P., Corruble, E., Fève, B., Colle, R., Ripoll, L., Walther, B., Boursier-Neyret, C., Werner, E., Becquemont, L., & Chanson, P. (2017). The human plasma-metabolome: Reference values in 800 French healthy volunteers; Impact of cholesterol, gender and age. PLoS ONE, 12(3), 1–17. 10.1371/journal.pone.0173615

Trachootham, D., Alexandre, J., & Huang, P. (2009). Targeting cancer cells by ROS-mediated mechanisms: A radical therapeutic approach? Nature Reviews Drug Discovery, 8(7), 579–591. 10.1038/nrd2803

Ubellacker, J. M., Tasdogan, A., Ramesh, V., Shen, B., Mitchell, E. C., Martin-Sandoval, M. S., Gu, Z., McCormick, M. L., Durham, A. B., Spitz, D. R., Zhao, Z., Mathews, T. P., & Morrison, S. J. (2020). Lymph protects metastasizing melanoma cells from ferroptosis. Nature, 585(7823), 113–118. 10.1038/s41586-020-2623-z

Upadhyayula, P. S., Higgins, D. M., Mela, A., Banu, M., Dovas, A., Zandkarimi, F., Patel, P., Mahajan, A., Humala, N., Nguyen, T. T. T., Chaudhary, K. R., Liao, L., Argenziano, M., Sudhakar, T., Sperring, C. P., Shapiro, B. L., Ahmed, E. R., Kinslow, C., Ye, L. F., … Canoll, P. (2023). Dietary restriction of cysteine and methionine sensitizes gliomas to ferroptosis and induces alterations in energetic metabolism. Nature Communications, 14(1). 10.1038/s41467-023-36630-w

Vande Voorde, J., Ackermann, T., Pfetzer, N., Sumpton, D., Mackay, G., Kalna, G., Nixon, C., Blyth, K., Gottlieb, E., & Tardito, S. (2019). Improving the metabolic fidelity of cancer models with a physiological cell culture medium. Science Advances, 5(1), eaau7314. 10.1126/sciadv.aau7314

Vlachogiannis, G., Hedayat, S., Vatsiou, A., Jamin, Y., Fernández-Mateos, J., Khan, K., Lampis, A., Eason, K., Huntingford, I., Burke, R., Rata, M., Koh, D., Tunariu, N., Collins, D., Hulkki-Wilson, S., Ragulan, C., Spiteri, I., Moorcraft, S. Y., Chau, I., … Valeri, N. (2018). Patient-derived organoids model treatment response of metastatic gastrointestinal cancers. Science, 359(6378), 920–926. 10.1126/science.aao2774

Vollmer, S., Kappler, V., Kaczor, J., Flügel, D., Rolvering, C., Kato, N., Kietzmann, T., Behrmann, I., & Haan, C. (2009). Hypoxia-inducible factor 1α is up-regulated by oncostatin M and participates in oncostatin M signaling. Hepatology, 50(1), 253–260. 10.1002/hep.22928

Wang, X., Zhao, X., Chou, J., Yu, J., Yang, T., Liu, L., & Zhang, F. (2018). Taurine, glutamic acid and ethylmalonic acid as important metabolites for detecting human breast cancer based on the targeted metabolomics. Cancer Biomarkers, 23(2), 255–268. 10.3233/CBM-181500

Whiteman, D. C., Green, A. C., & Olsen, C. M. (2016). The Growing Burden of Invasive Melanoma: Projections of Incidence Rates and Numbers of New Cases in Six Susceptible Populations through 2031. Journal of Investigative Dermatology, 136(6), 1161–1171. 10.1016/j.jid.2016.01.035

Winder, M., & Virós, A. (2018). Mechanisms of drug resistance in Melanoma. In Handbook of Experimental Pharmacology (Vol. 249). 10.1007/164_2017_17

