## Supplemental figures for "Ferroptosis susceptibility of melanoma cells: dependence on cell-type, acquired drug resistance, and medium composition"

### Table S1

## A

| Solution 1: | Conc.(μM) | Provider | order number |
|---|---|---|---|
| L-Histidine | **110** | Roth | 1696.1 |
| L-Isoleucine | **77** | Roth | 1698.1 |
| L-Leucine | **132** | Roth | 1699.1 |
| L-Lysine (monohydrate) | **210** | Roth | 6829.1 |
| L-Methionine | **25** | Roth | 1702.1 |
| L-Phenylalanine | **75** | Roth | 1709.1 |
| L-Threonine | **142** | Roth | 1738.1 |
| L-Tryptophan | **60** | Roth | 1739.1 |
| L-Valine | **230** | Roth | 1742.1 |
| L-Citrulline | **37** | Sigma/Merck | C7629 |
| L-Ornitine (monohydrochloride) | **74** | Roth | 1703.1 |

| Solution 2: | Conc.(μM) | Provider | order number |
|---|---|---|---|
| L-Alanine | **435** | Roth | 3076.1 |
| L-Arginine | **100** | Roth | 1655.1 |
| L-Proline | **210** | Roth | 1713.1 |
| L-Tyrosine | **76** | Roth | 1741.1 |

| Solution 3: | Conc.(μM) | Provider | order number |
|---|---|---|---|
| L-Asparagine (monohydrate) | **50** | Roth | KK37.1 |
| L-Aspartic acid | **13** | Roth | 1690.1 |
| L-Glutamic acid | **90** | Roth | 1743.1 |
| Glycine | **295** | Roth | HN07.1 |
| L-Serine | **140** | Roth | 1714.1 |
| L-Cystine | **67** | Roth | 1695.1 |
| L-Cysteine | **32** | Roth | 1693.1 |

| Solution 4: | Conc.(μM) | Provider | order number |
|---|---|---|---|
| a-Aminobutyrate (L-2-aminobutyric acid) | **26** | Alpha Aesar | L13984.03 |
| L-Pyroglutamate | **100** | Sigma/Merck | 83160 |
| 4-Hydroxy-L-proline (trans) | **13** | Alpha Aesar | A11851.09 |
| L-Acetyl glycine | **85** | Sigma/Merck | A16300 |
| Glutathione (reduced) | **31** | Roth | 6832.1 |
| Taurine | **104** | Roth | 4721.1 |
| N-Trimethylglycine (betaine) | **74** | Sigma/Merck | 61962 |
| Acetone | **68** | Sigma/Merck | 650501 |
| Acetyl carnitine (hydrochloride) | **6** | Sigma/Merck | A6706 |
| L-Carnitine | **40** | Roth | A17618.09 |
| Creatine | **37** | Sigma/Merck | C0780 |
| Creatinine | **80** | Alpha Aesar | B23097.09 |
| Glycerol | **95** | Sigma/Merck | G5516 |
| 2-Hydroxybutyrate (Sodium salt) | **55** | Sigma/Merck | 220116 |
| 3-Hydroxybutyrate (Sodium salt) | **103** | Sigma/Merck | 54965 |
| Hypoxanthine | **14** | Alpha Aesar | A11481.06 |
| Methyl acetoacetate | **40** | Sigma/Merck | 537365 |
| Urea | **4000** | Roth | 2317.1 |

| Solution 5: | Conc.(μM) | Provider | order number |
|---|---|---|---|
| Succinate | **21** | Sigma/Merck | S2378 |
| Lactate (Sodium salt) | **1000** | Sigma/Merck | 71718 |
| Formate (Sodium salt) | **60** | Alpha Aesar | A17813.30 |
| Pyruvate (Sodium salt) | **93** | Alpha Aesar | A11148.18 |
| Citrate (Sodium salt, 5.5 hydrate) | **115** | Sigma/Merck | C8532 |
| Acetate (Sodium salt) | **40** | Sigma/Merck | S2889 |

| Solution 6: | Conc.(μM) | Provider | order number |
|---|---|---|---|
| Ammonium chloride | **50** | Fluka | 9700 |
| Iron sulfate (heptahydrate) | **0.33** | Merck | 103965.05 |
| Sodium nitrate | **0.016** | Sigma/Merck | S8170 |
| Zinc chloride | **0.51** | Sigma/Merck | 1.0881 |

| Solution 7: | Conc.(μM) | Provider | order number |
|---|---|---|---|
| Sodium selenite | **0.02911** | Sigma/Merck | 214485 |

| Solution 8: | Conc.(μM) | Provider | order number |
|---|---|---|---|
| Urate (uric acid sodium salt) | **270** | Sigma/Merck | U2875 |

| Solution9: | Conc.(μM) | Provider | order number |
|---|---|---|---|
| L-Glutamine | **513** | Roth | HN08.1 |

| Vitamin Mix: | Conc.(μM) | Provider | order number |
|---|---|---|---|
| D-Biotin | **4.10** | Sigma/Merck | B6891 |
| Choline chloride | **7.10** | | |
| Folate | **2.30** | | |
| myo-Inositol | **11.10** | | |
| Niacinamide | **8.20** | | |
| D-Pantothenic acid hemicalcium | **4.20** | | |
| Pyridoxine | **4.90** | | |
| Riboflavin | **0.30** | | |
| Thiamine | **3.00** | | |
| Ascorbic acid | **68.0** | Sigma/Merck | A4034 |
| Vitamin B12 | **0.015** | Roth | T915.1 |

| EBSS: | Conc.(μM) | Provider | order number |
|---|---|---|---|
| Calcium chloride | **1622** | Gibco | 24010043 |
| Magnesium sulfate (heptahydrate) | **732** | | |
| Potassium chloride | **4829** | | |
| Sodium bicarbonate | **23569** | | |
| Sodium chloride | **104723** | | |
| Sodium phosphate monobasic (hydrate) | **913** | | |
| D-Glucose | **4996** | | |
| Phenol red | **2.54** | | |

## B

| B27 supplement | Conc. B18 (μM) |
|---|---|
| Bovine serum albumin, fatty acid free | 37 |
| Bovine holo transferrin | 0.062 |
| Catalase | 0.01 |
| Superoxide dismutase | 0.077 |
| Insulin | 0.6 |
| Linoleic acid | 3.5 |
| Linolenic acid | 3.5 |
| T3 | 0.026 |
| Progesterone | 0.020 |
| Corticosterone | 0.058 |
| Retinol acetate | 0.2 |
| D,L-alpha tocopherol | 2.3 |
| D,L-alpha tocopherol acetate | 2.1 |
| Biotin | 0.4 |
| D-Galactose | 83 |
| Ethanolamine | 16 |
| L-Carnitine | 12 |
| Putrescine | 183 |
| Selenite | 0.083 |
| Glutathione reduced | 3.2 |

### Table S2

The numbers indicated in the table are concentrations in $\mu M$

| | Physiologic media | | | Standard cell culture media | | | |
|---|---|---|---|---|---|---|---|
| Proteinogenic amino acids | MPM | HPLM | Plasmax | RPMI1640 | MEM | DMEM | IMDM |
| Glutamine | 513 | 550 | 650 | 2055 | 2000 | 4000 | 4000 |
| Alanine | 435 | 430 | 510 | | | | 281 |
| Glycine | 295 | 300 | 330 | 133 | | 400 | 400 |
| Valine | 230 | 220 | 230 | 171 | 393 | 803 | 803 |
| Lysine | 210 | 200 | 220 | 219 | 399 | 798 | 798 |
| Proline | 210 | 200 | 360 | 174 | | | 348 |
| Threonine | 142 | 140 | 240 | 168 | 403 | 798 | 798 |
| Serine | 140 | 150 | 140 | 286 | | 400 | 400 |
| Leucine | 132 | 160 | 170 | 382 | 397 | 802 | 802 |
| Histidine | 110 | 110 | 120 | 97 | 200 | 200 | 200 |
| Arginine | 100 | 110 | 64 | 1150 | 597 | 398 | 399 |
| Glutamic acid | 90 | 80 | 98 | 136 | | | 510 |
| Isoleucine | 77 | 70 | 140 | 382 | 397 | 802 | 802 |
| Tyrosine | 76 | 80 | 74 | 111 | 199 | 399 | 462 |
| Phenylalanine | 75 | 80 | 68 | 91 | 194 | 400 | 400 |
| Cystine | 67 | 100 | 65 | 207 | 99 | 201 | 292 |
| Tryptophan | 60 | 60 | 78 | 25 | 49 | 78 | 78 |
| Asparagine | 50 | 50 | 41 | 379 | | | 189 |
| Cysteine | 32 | 40 | 33 | | | | |
| Methionine | 25 | 30 | 30 | 101 | 101 | 201 | 201 |
| Aspartic acid | 13 | 20 | 6 | 150 | | | 226 |

| Other metabolites | MPM | HPLM | Plasmax | RPMI1640 | MEM | DMEM | IMDM |
|---|---|---|---|---|---|---|---|
| D-Glucose | 4996 | 5000 | 5551 | 11111 | 5556 | 25000 | 25000 |
| Urea | 4000 | 5000 | 3000 | | | | |
| Lactate | 1000 | 1600 | 500 | | | | |
| Urate | 270 | 350 | 270 | | | | |
| Citrate | 115 | 130 | 114 | | | | |
| Taurine | 104 | 90 | 130 | | | | |
| 3-Hydroxybutyrate | 103 | 50 | 77 | | | | |
| L-Pyroglutamate2 | 100 | | 20 | | | | |
| Glycerol | 95 | 120 | 82 | | | | |
| Pyruvate | 93 | 50 | 100 | | | 1000 | 1000 |
| L-Acetylglycine | 85 | 90 | 70 | | | | |
| Creatinine | 80 | 75 | 74 | | | | |
| L-Ornitine | 74 | 70 | 80 | | | | |
| N-Trimethylglycine | 74 | 70 | 72 | | | | |
| Acetone | 68 | 60 | 55 | | | | |
| Formate | 60 | 50 | 33 | | | | |
| 2-Hydroxybutyrate | 55 | 50 | 31 | | | | |
| L-Carnitine | 40 | 40 | 46 | | | | |
| Methylacetoacetate | 40 | | 41 | | | | |
| Acetate | 40 | 40 | 42 | | | | |
| L-Citrulline | 37 | 40 | 55 | | | | |
| Creatine | 37 | 40 | 37 | | | | |
| Glutathione | 31 | 25 | 37 | 3.3 | | | |
| $\alpha$-Aminobutyrate | 26 | 20 | 41 | | | | |
| Succinate | 21 | 20 | 23 | | | | |
| Hypoxanthine | 14 | 10 | 5 | | | | |
| 4-Hydroxy-L-proline | 13 | 20 | 13 | | | | |
| Acetyl carnitine | 6 | | 5 | | | | |
| L-Homocysteine | | | 9 | | | | |
| L-Carnosine | | | 6 | | | | |
| Malonate | | 10 | | | | | |
| Uracil | | | 2 | | | | |
| Uridine | | | 3 | | | | |
| D-Fructose | | 40 | | | | | |
| D-Galactose | | 60 | | | | | |

| Vitamins | MPM | HPLM | Plasmax | RPMI1640 | MEM | DMEM | IMDM |
|---|---|---|---|---|---|---|---|
| Thiamine | 3.0 | 3.0 | 3.0 | 3.0 | 3.0 | 11.2 | 11.9 |
| Riboflavin | 0.3 | 0.5 | 0.3 | 0.5 | 0.3 | 1.1 | 1.1 |
| Nicotin amide | 8.2 | 8.2 | 8.2 | 8.2 | 8.2 | 32.8 | 32.8 |
| Choline chloride | 7.1 | 21.5 | 7.1 | 21.4 | 7.1 | 28.6 | 28.6 |
| D-Pantothenic acid hemicalcium | 4.20 | 1.05 | 4.20 | 0.52 | 2.10 | 8.40 | 8.40 |
| Pyridoxal | | | | | 4.9 | | 19.6 |
| Pyridoxine | 4.9 | 4.9 | 4.9 | 4.9 | | 19.6 | |
| D-Biotin | 4.10 | 0.80 | 4.10 | 0.82 | | | 0.53 |
| myo-inositol | 11.1 | 194.3 | 11.1 | 194.4 | 11.1 | 40.0 | 40.0 |
| Folate | 2.30 | 2.30 | 2.30 | 2.27 | 2.30 | 9.10 | 9.10 |
| Vitamin B12 | 0.0150 | 0.0037 | 0.0050 | 0.0037 | | | 0.0096 |
| Para-aminobenzoic acid | | 7.3 | | 7.3 | | | |
| Ascorbic acid | 68.0 | | 62.0 | | | | |

| Inorganic salts | MPM | HPLM | Plasmax | RPMI1640 | MEM | DMEM | IMDM |
|---|---|---|---|---|---|---|---|
| Calcium chloride | 1622 | 2350 | 1802 | | 1802 | 1802 | 1487 |
| Calcium nitrate | | 40 | | 425 | | | |
| Magnesium chloride | | 480 | | 602 | | | |
| Magnesium sulfate | 732 | 350 | 813 | 407 | 814 | 814 | 814 |
| Potassium chloride | 4829 | 4100 | 5365 | 5333 | 5333 | 5333 | 4400 |
| Potassium nitrate | | | | | | | 1 |
| Sodium bicarbonate | 23569 | 24000 | 26187 | 23810 | 26192 | 44048 | 36000 |
| Sodium chloride | 104723 | 105000 | 116359 | 103448 | 117241 | 110345 | 77672 |
| Sodium phosphate monobasic | 913 | 870 | 1014 | | 1014 | 906 | 906 |
| Sodium phosphate dibasic | | | | 5634 | | | |
| Ammonium chloride | 50 | 40 | 50 | | | | |
| Iron nitrate | | | 0.012 | | | | |
| Iron sulfate | 0.33 | | 1.04 | | | | |
| Iron chloride | | | | | | | 2 |
| Ferric nitrate | | | | | | 0.2475 | |
| Sodium nitrate | 0.016 | | 0.032 | | | | |
| Zinc chloride | 0.51 | | 1.5 | | | | |
| Zinc sulfate | | | | | | | 0.490 |
| Sodium selenite | 0.029 | | 0.029 | | | | 0.098 |
| Cupric sulfate | | | 0.0052 | | | | |
| Manganese chloride | | | 0.0002 | | | | |
| Ammonium metavanadate | | | 0.0026 | | | | |

### Figure S1

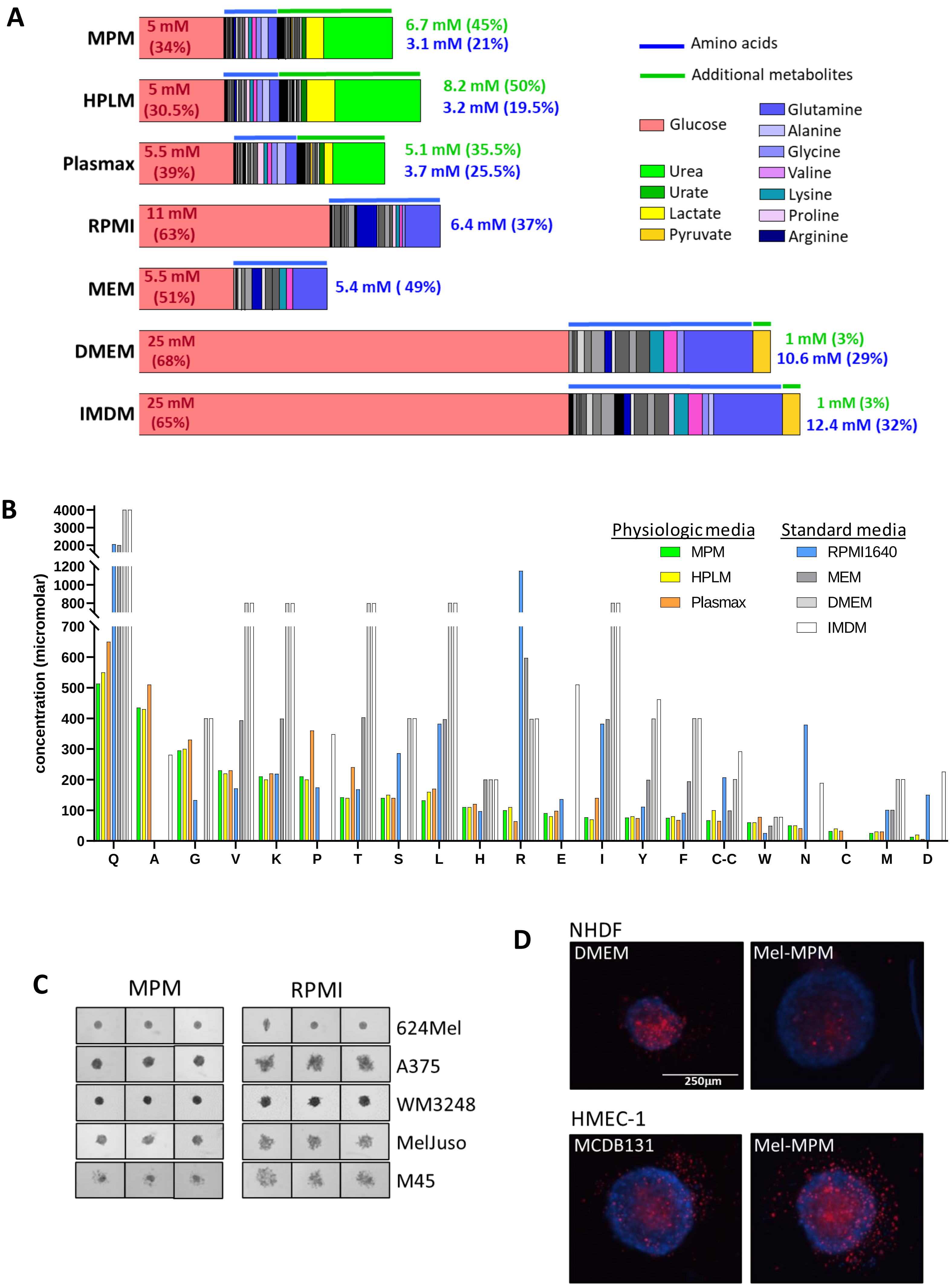
A
MPM
5 mM (34%)
6.7 mM (45%)
3.1 mM (21%)
HPLM
5 mM (30.5%)
8.2 mM (50%)
3.2 mM (19.5%)
Plasmax
5.5 mM (39%)
5.1 mM (35.5%)
3.7 mM (25.5%)
RPMI
11 mM (63%)
6.4 mM (37%)
MEM
5.5 mM (51%)
5.4 mM ( 49%)
DMEM
25 mM (68%)
1 mM (3%)
10.6 mM (29%)
IMDM
25 mM (65%)
1 mM (3%)
12.4 mM (32%)
Amino acids
Additional metabolites
Glucose
Urea
Urate
Lactate
Pyruvate
Glutamine
Alanine
Glycine
Valine
Lysine
Proline
Arginine
B
concentration (micromolar)
4000
3000
2000
1200
1000
800
700
600
500
400
300
200
100
0
Q A G V K P T S L H R E I Y F C-C W N C M D
Physiologic media
MPM
HPLM
Plasmax
Standard media
RPMI1640
MEM
DMEM
IMDM
C
MPM
RPMI
624Mel
A375
WM3248
MelJuso
M45
D
NHDF
DMEM
Mel-MPM
250µm
HMEC-1
MCDB131
Mel-MPM

**Figure S2:**

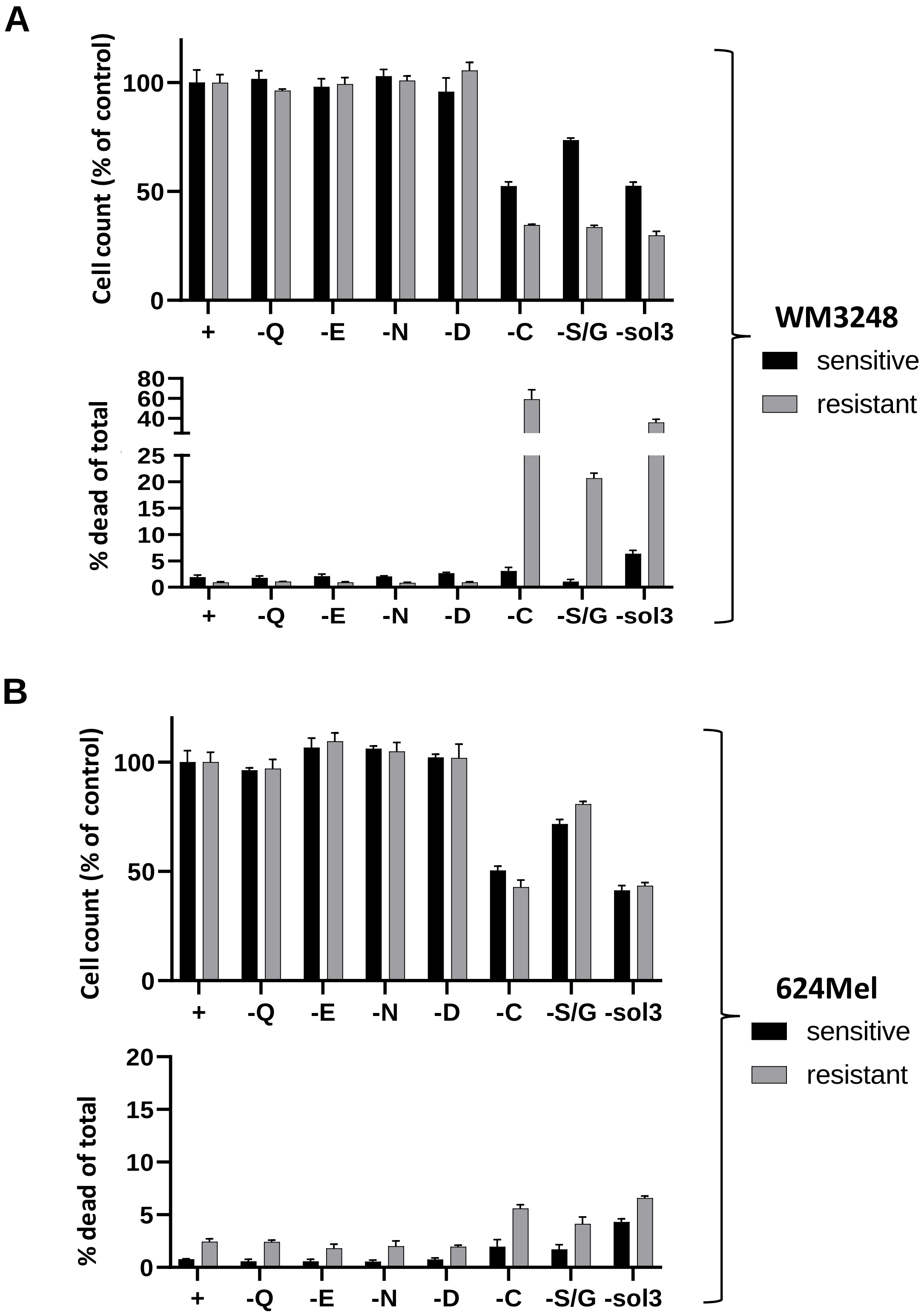
A
Cell count (% of control)
100
50
0
+
-Q
-E
-N
-D
-C
-S/G
-sol3
% dead of total
80
60
40
25
20
15
10
5
0
WM3248
sensitive
resistant
B
624Mel
sensitive
resistant

**Figure S3:**

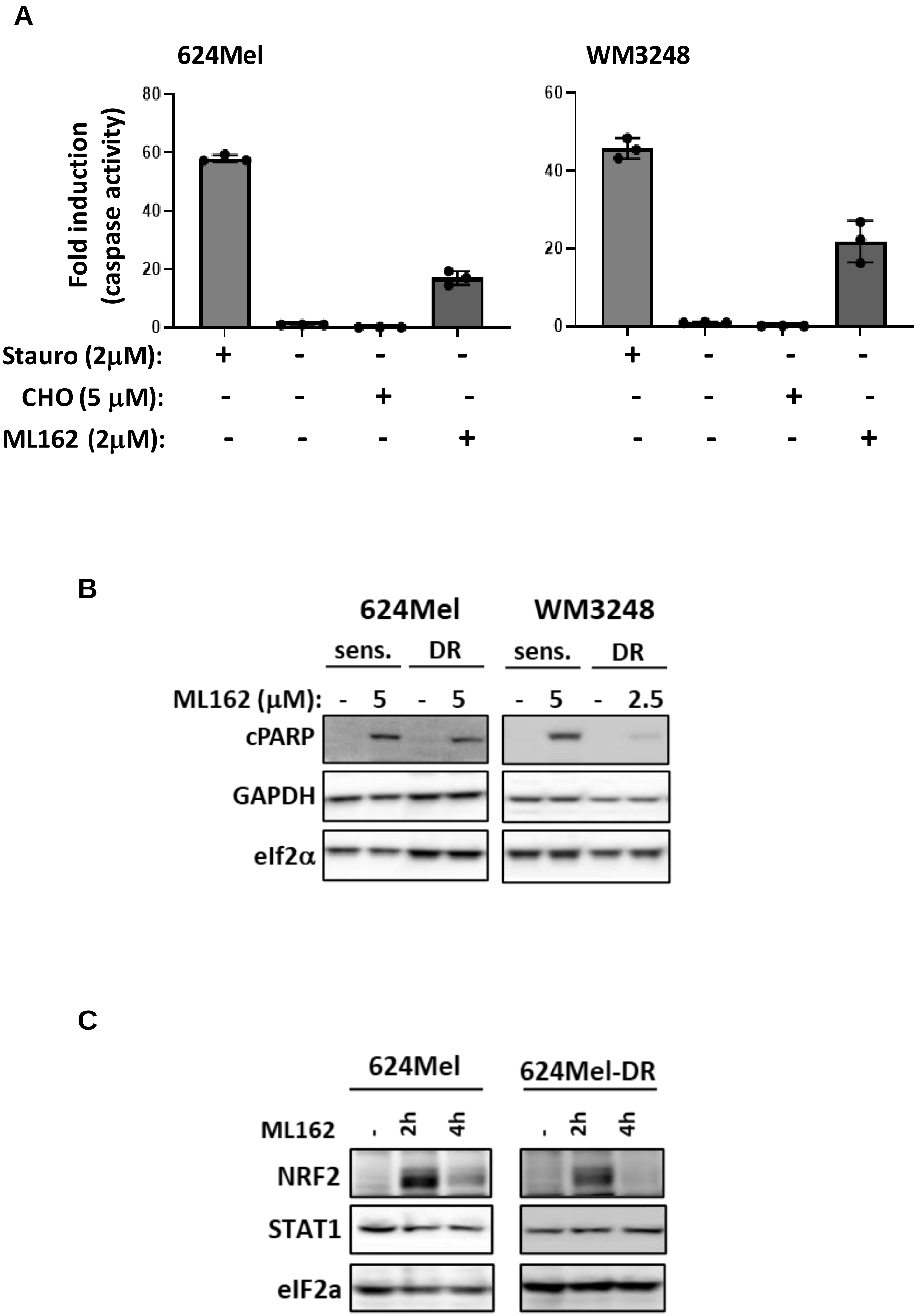
A
624Mel
WM3248
Fold induction (caspase activity)
80
60
40
20
0
Stauro (2μM): + - - -
CHO (5 μM): - - + -
ML162 (2μM): - - - +
B
624Mel
WM3248
sens.
DR
ML162 (μM): - 5 - 5 - 5 - 2.5
cPARP
GAPDH
eIf2α
C
624Mel
624Mel-DR
ML162
-
2h
4h
NRF2
STAT1
eIF2a

### Figure S4

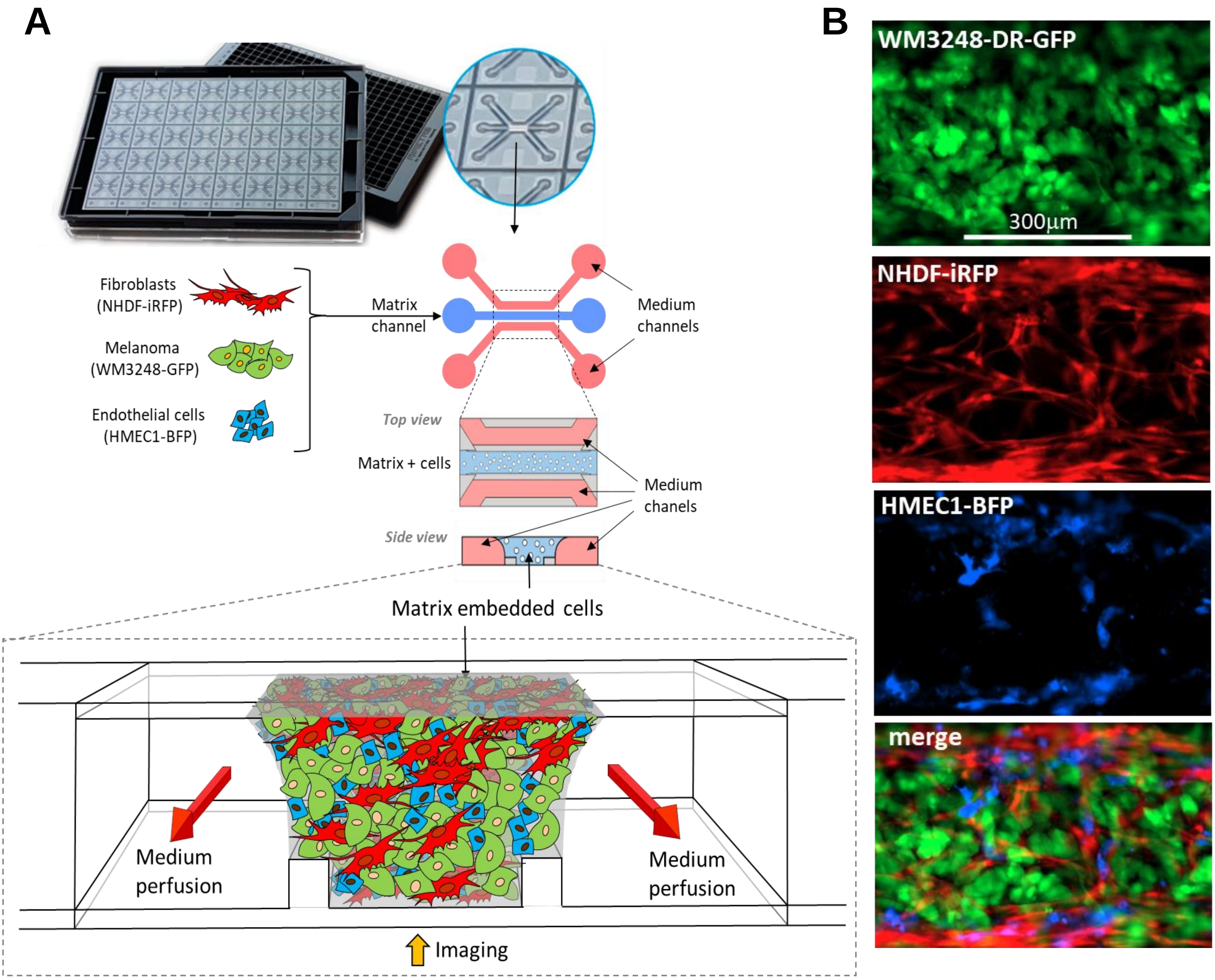
A
Fibroblasts
(NHDF-iRFP)
Melanoma
(WM3248-GFP)
Endothelial cells
(HMEC1-BFP)
Matrix
channel
Medium
channels
Top view
Matrix + cells
Medium
chanels
Side view
Matrix embedded cells
Medium
perfusion
Medium
perfusion
Imaging
B
WM3248-DR-GFP
300μm
NHDF-iRFP
HMEC1-BFP
merge

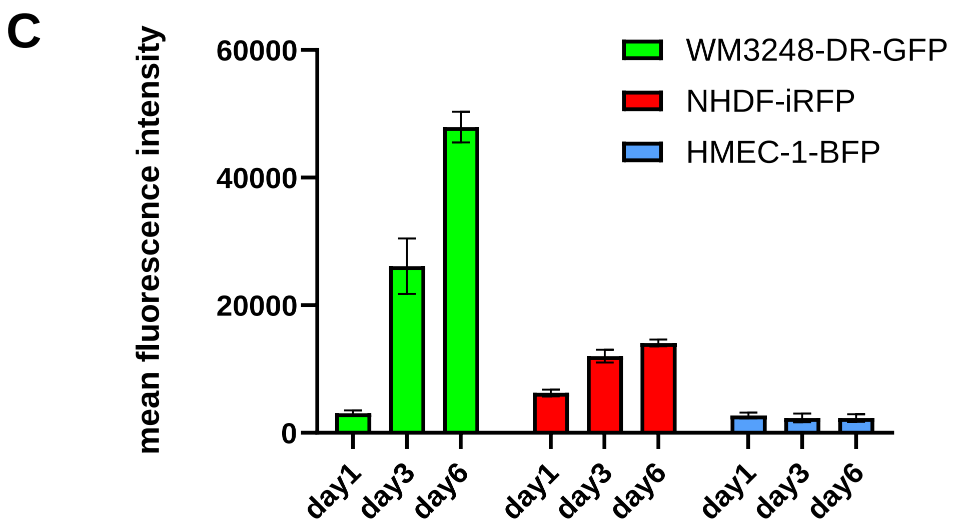
C
mean fluorescence intensity
60000
40000
20000
0
WM3248-DR-GFP
NHDF-iRFP
HMEC-1-BFP
day1
day3
day6
day1
day3
day6
day1
day3
day6

#### Supplementary Figures

**Table S1. Composition of the different solutions making up Modular Physiologic Medium (MPM) and Mel-MPM.** A) The concentration, provider, and ordering number for each constituent of MPM is shown. B) Composition of B27 (Brewer et al., 1993) and concentrations of B18 (Brewer & Cotman, 1989). Note that for B18 the concentrations of the components are published, while the exact concentrations of B27 are not published. However, the identity of the compounds in B27 is published and identical to B18.

**Table S2. Comparison of the composition of different physiologic and standard cell culture media**

The concentrations of the components (in μM) for the modular physiologic medium MPM used in this study, Plasmax (Vande Voorde et al., 2019) and HPLM (Cantor et al., 2017), and for the standard cell culture media RPMI1640, MEM, DMEM, and IMDM are indicated.

**Figure S1. Comparative presentation of main nutrient contents (glucose, amino acids and other metabolites) of physiologic media (HPLM, Plasmax and MPM) and the classical culture media RPMI1640, MEM, DMEM, and IMDM.** A) Bar charts representing the nutrient composition of the media. The total concentration and the percentage in relation to the total nutrient content is indicated, for glucose (red), amino acids (blue lines and writing), and other metabolites (green lines and writing). Specific metabolites are highlighted by colors. The number of supra-physiologically concentrated amino acids is indicated in blue italics. The individual amino acid concentrations can be identified in B. B) The concentrations of all amino acids are shown for the physiologic media HPLM, Plasmax and MPM and for the standard media RPMI1640, MEM, DMEM and IMDM. The graph highlights the similarity of the physiologic media and the huge differences of concentrations of amino acids found in standard cell culture media. Note: In RPMI, the medium in which many melanoma cells grow, 9 amino acids are present at concentration above physiological concentrations; alanine, the second most abundant amino acid in plasma, is absent. In DMEM, the medium in which some of the melanoma cell lines and the NHDF grow, the concentrations of 13 amino acids are drastically increased, 5 amino acids are lacking. C) Melanoma cell lines were tested for their sphere forming capacity in MPM compared to RPMI. D) Microscopic picture of 3D spheroids of NHDF (upper panel) and HMEC1 (lower panel) cells in their standard media (DMEM and MCDB131, respectively) compared to Mel-MPM. Nuclei are stained in blue with Hoechst 33342. Dead cells are stained in red using PI.

**Figure S2. Investigation of the consequences of total withdrawal of individual non-essential amino acids (NEAA) in drug-resistant versus drug-naïve cells using 2D growth assays.** 2D growth assays were performed with the drug-sensitive and -resistant WM3248 (A) and 624Mel cells (B) in MPM lacking specifically one NEAA (the one letter code is used to indicate the omitted amino acid). Cystine/cysteine (-C) and serine/glycine ("-S/G") were depleted in combination, while the other amino acids were depleted individually; -sol3: solution 3 was omitted during MPM preparation. + represents MPM with all components. Numbers of living or dead cells were determined by applying a live nuclear cell stain (Hoechst 33342) and a dead cell stain (Sytox orange) using a Cytation 5 instrument and the Gen5 software. Graphs showing % of living (compared to control cells in MPM) versus % of dead cells are shown side by side.

**Figure S3.** A) Caspase activity assays of 624Mel and WM3248 cells treated with ML162 for 16 hours, while being cultured in MPM. Staurosporine (Stauro) was used as positive control. The caspase inhibitor Ac-DEVD-CHO (CHO) was used to give an idea about the background apoptosis in the different cultures, which was negligible compared to the compound-induced one. B) Western blots of total cell lysates of 624Mel, 624Mel-DR, WM3248, and WM3248-DR cells treated with ML162 for 16 hours in MPM. The blots were detected with a specific c-PARP antibody to show Caspase 3-dependent PARP cleavage occurring during apoptosis. GAPDH and eIF2$\alpha$ detections were performed as loading controls. C) Lysates of 624Mel and 624Mel-DR cells treated or not with ML162 (for 2 or 4 hours) were subjected to SDS-PAGE and Western blot analysis. Protein levels of NRF2, upregulated by ML162 treatment are shown. STAT1 and eIF2$\alpha$ levels are shown as loading controls.

**Figure S4. The OrganoPlates™ microchannel system used to implement the complex matrix embedded 3D multi-cell type system.** A) Scheme of a 3-channel 40 OrganoPlate™ and the matrix-embedded cells each expressing different fluorescent proteins (BFP, GFP, iRFP). The matrix-embedded cells were perfused on both sides by drug-containing Mel-MPM medium. Scheme adapted from Kramer et al., 2019. B) 10,000 WM3248-DR-GFP, 5,000 NHDF-iRFP, and 5,000 HMEC-1BFP per µL Geltrex were seeded in the OrganoPlate™. After 2 days, the medium was replaced, and the cells were left to grow for 4 days. The cells were imaged using the Cytation 10 instrument using a 10x objective and the LED/Filter cubes for GFP, Cy5.5, and DAPI. Live cells are detected by the fact that they express their respective fluorescent protein. C) The mean fluorescence intensity of the different fluorophores expressed in the 3 cell types is shown over the course of 6 days to illustrate the growth behaviour of the different cell types in the matrix.
